# Nucleoside-Modified mRNA Encoding Alpha-Galactosidase A Reverses Fabry Disease Phenotypes in Human IPSC-Derived Cardiomyocytes

**DOI:** 10.64898/2025.12.03.692072

**Authors:** Malte Juchem, Sedef Ersoy, Lea Oehlsen, Jia Li Ye, Natalie Weber, Ke Xiao, Christopher Jahn, Reto Eggenschwiler, Tobias Cantz, Julia Beimdiek, Falk F. R. Buettner, Theresia Kraft, Jan Hegermann, Christian Bär, Jeannine Hoepfner, Thomas Thum

## Abstract

The lysosomal storage disorder Fabry disease results from α-galactosidase A deficiency, leading to excessive glycosphingolipid substrate accumulation, primarily globotriaosylceramide (Gb3). While the underlying molecular mechanisms remain elusive, multi-systemic complications ultimately culminate in premature death, with heart failure being the leading cause of death. Current treatment options fail to treat Fabry disease adequately and only delay its progression. Preclinical studies on an alternative approach, systemic delivery of nucleoside-modified *GLA* mRNA (modGLA), suggest improved effectiveness over existing therapies in reducing glycosphingolipid levels in the heart.

It remains unclear whether modGLA can rescue Fabry cardiomyopathy phenotypes at the cellular level, which are not faithfully recapitulated in current animal models. To address this, we investigated characteristic phenotypes in two new models of Fabry cardiomyopathy utilizing human iPSC-derived cardiomyocytes in transcriptomic and functional analyses. These human Fabry disease cardiomyocytes displayed broad transcriptional dysregulation, apoptosis, mitochondrial dysfunction, impaired reactive oxygen species handling, as well as enhanced decay parameters of calcium transients. Mechanistically, we identified hyperphosphorylated phospholamban as a major player in this calcium dysregulation. Strikingly, modGLA therapy of Fabry cardiomyocytes restored α-galactosidase A enzyme activity, reduced glycosphingolipid deposition, and normalized the observed molecular alterations, supporting modGLA therapy as a promising strategy for the treatment of Fabry disease.

## Introduction

Fabry disease (OMIM #301500) is an X-linked lysosomal storage disorder that results from deficiency of the lysosomal enzyme α-galactosidase A, encoded by pathogenic *GLA* variants^[1]^. Consequently, the impaired metabolism of its glycosphingolipid substrates, primarily globotriaosylceramide (Gb3)^[2]^, causes their build-up throughout patients’ tissues. This glycosphingolipid accumulation is discussed to modulate secondary biochemical effects including membrane lipid composition, protein trafficking and mitochondrial function^[3]^, which then progressively manifest as a multisystemic disorder, most severely affecting the cerebral, renal, and cardiovascular system. The primary cause of death are cardiovascular complications both in affected men and women^[4]^. Typically observed cardiac phenotypes include left ventricular hypertrophy, arrhythmias, myocardial fibrosis and eventually heart failure^[5]^. On cellular and tissue level oxidative stress and apoptosis^[6]^, mitochondrial dysfunction and consequently impaired cardiac energy metabolism as well as inflammation,^[7]^ have been reported.

Unfortunately, Fabry disease remains inadequately addressed by the existing treatment options. Although available therapeutic options for Fabry disease have been demonstrated to temporarily stall disease progression, patients continue to experience a considerable disease burden that culminates in premature death^[8,9]^. While enzyme replacement therapy (ERT) appears to slow disease progression it fails to remove Gb3 from affected cardiomyocytes or to ameliorate heart dysfunction even after follow-up of 20 years^[10,11]^. The only approved alternative is based on pharmacological chaperones, which are only available to a subset of patients with amenable mutations^[12]^.

The Fabry pathomechanism remains insufficiently understood. Unravelling the underlying molecular mechanisms might allow the development of novel treatment approaches. However, the established animal models including *GLA*-KO mice^[13,14]^ and rats^[15]^ fail to mirror the consequences of *GLA* deficiency observed in Fabry patients, particularly the cardiac manifestation. Consequently, alternative models are required to study disease mechanisms and test the effectiveness of novel treatments in a relevant and translational system. To approach this, we aimed to establish such an alternative, a general and patient-independent *in vitro* Fabry disease model by knockout of *GLA* in hiPSCs derived from a healthy donor. Subsequently, we utilized this model system to investigate the effectiveness of the promising novel candidate-treatment ‘nucleoside-modified *GLA* mRNA (modGLA)’ in ameliorating Fabry-typical cellular disease phenotypes.

The SARS-CoV 2 pandemic illustrated the immense potential of RNA-based drugs. Nucleoside modifications may have partly facilitated their wide-spread use^[16]^. In the context of Fabry disease, preclinical studies using *GLA* mRNA have shown an improved therapeutic efficacy in the hearts and kidneys of *GLA*-KO mice, specifically regarding a reduction of Gb3 and lyso-Gb3 levels, when compared to ERT alone, ERT and chaperone therapy combined, or other gene therapy approaches^[17]^. Yet, established animal models fail to recapitulate the complex cellular pathology of Fabry disease. *In vitro* data for modGLA therapy in human Fabry cardiomyocytes remain limited to reduction of Gb3 deposition and partial normalization of lysosomal protein levels^[18]^. Here, we investigate whether nucleoside-modified *GLA* mRNA can rescue disease-relevant cellular phenotypes beyond substrate accumulation and provide mechanistic insight into cardiomyocyte dysfunction in Fabry disease.

## Results

### Loss of *GLA* results in accumulation of the α-galactosidase A substrate globotriaosylceramide in hiPSC-derived cardiomyocytes

To investigate the specific effects of functional *GLA* loss underlying the cardiac Fabry disease manifestation, while minimizing confounding genetic factors putatively present in patient-derived hiPSC-based *in vitro* disease models, we previously generated *GLA*-knockout (KO) hiPSCs from a healthy donor-derived *GLA* wild type (WT) parental line using CRISPR/Cas9^[19,20]^. In the present study, both *GLA*-KO and parental WT hiPSC lines readily differentiated into hiPSC-derived cardiomyocytes (CMs) as indicated by 95.36 (0.93)% and 94.63 (0.68)% cardiac troponin T-positive cells for WT and *GLA*-KO CMs, respectively (Fig. 1 B). As expected, *GLA*-KO CMs were α-galactosidase A enzyme deficient (Fig. 1 D) which resulted in progressive accumulation of its glycosphingolipid substrate Gb3 (Fig. 1 C). Since myofibrillolysis has been anecdotally reported in endomyocardial biopsies from Fabry patients^[21]^, we performed transmission electron microscopy to analyze sarcomere ultrastructure. However, no apparent differences between the models were observed in 65-day-old CMs (Fig. 1 E).

**Figure 1.**
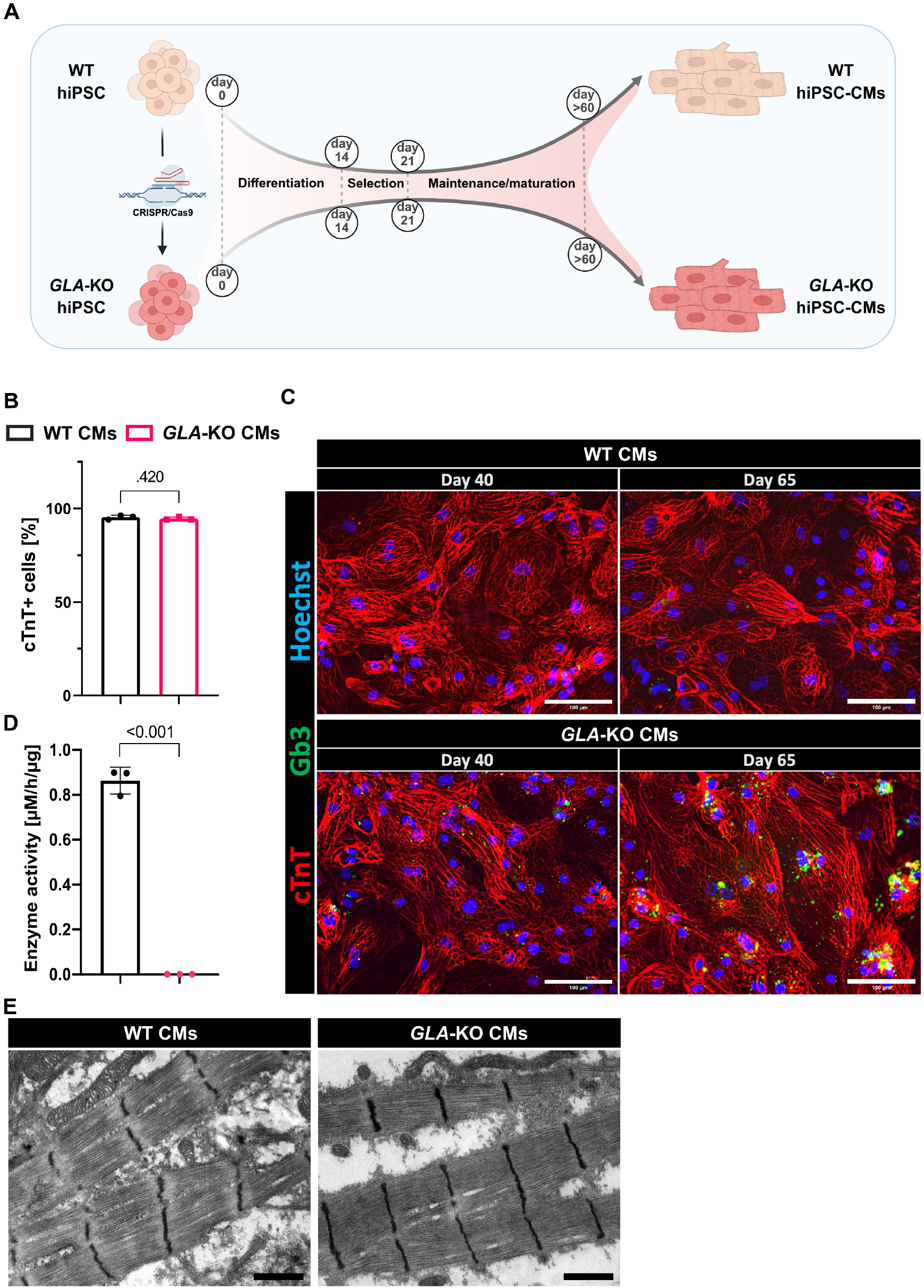
*GLA*-KO CMs are α-galactosidase A enzyme deficient and progressively accumulate Gb3. (A) Schematic for the generation of a patient-independent hiPSC-derived *in vitro* model for the cardiac manifestation of Fabry disease. Created in BioRender. Hoepfner, J. (2025) https://BioRender.com/kouwaxh. (B) Cardiac differentiation efficiency as analyzed by flow cytometry for cardiac troponin T (cTnT) positive cells; (n = 3 independent differentiations, mean (SD), unpaired t-test, α = 0.05). (C) Progressive Gb3 accumulation in cTnT-expressing CMs analyzed in 40- and 65-day-old CMs by fluorescence microscopy. Scale bars indicate 100 µm. (D) The α-galactosidase A enzyme activity measured in cell lysates of 65-days-old CMs; (n = 3 independent differentiations, unpaired t-test, α = 0.05). (E) Ultrastructural analysis of CM sarcomeres using transmission electron microscopy. Scale bars indicate 1 µm.

### Transcriptome profiles resemble key phenotypes of Fabry disease

To ascertain the capacity of present *in vitro* disease model to replicate the cardiac manifestation of Fabry disease and elucidate its pathomechanism, we performed bulk-RNA sequencing of rRNA-depleted total RNA isolated from 65-day-old WT and *GLA*-KO CMs (Fig. 2 A). When subjecting the subset of 2397 differentially expressed genes (DEGs; base mean > 5; p < 0.05) to enrichment analysis (david.ncifcrf.gov^[22]^), these DEGs were particularly enriched in pathways previously associated with Fabry disease (Fig. 2 B). Specifically, pathways related to apoptosis^[23]^, mitochondrial energy metabolism^[24]^, and reactive oxygen species^[25]^ (Fig. 2 C) were deregulated. DEGs were enriched in terms related to the physiological and pathophysiological function of the heart, namely, “cardiac muscle contraction” and “diabetic cardiomyopathy”. In addition, several terms belonging to neurodegenerative diseases that involve cellular dysfunction and macromolecule accumulation including Huntington, Alzheimer, amyotrophic lateral sclerosis, and prion disease were identified. Of potential interest, several signaling pathways, specifically, Ras/MAPK, p53 and retrograde endocannabinoid signaling were significantly enriched for the identified DEGs.

**Figure 2.**
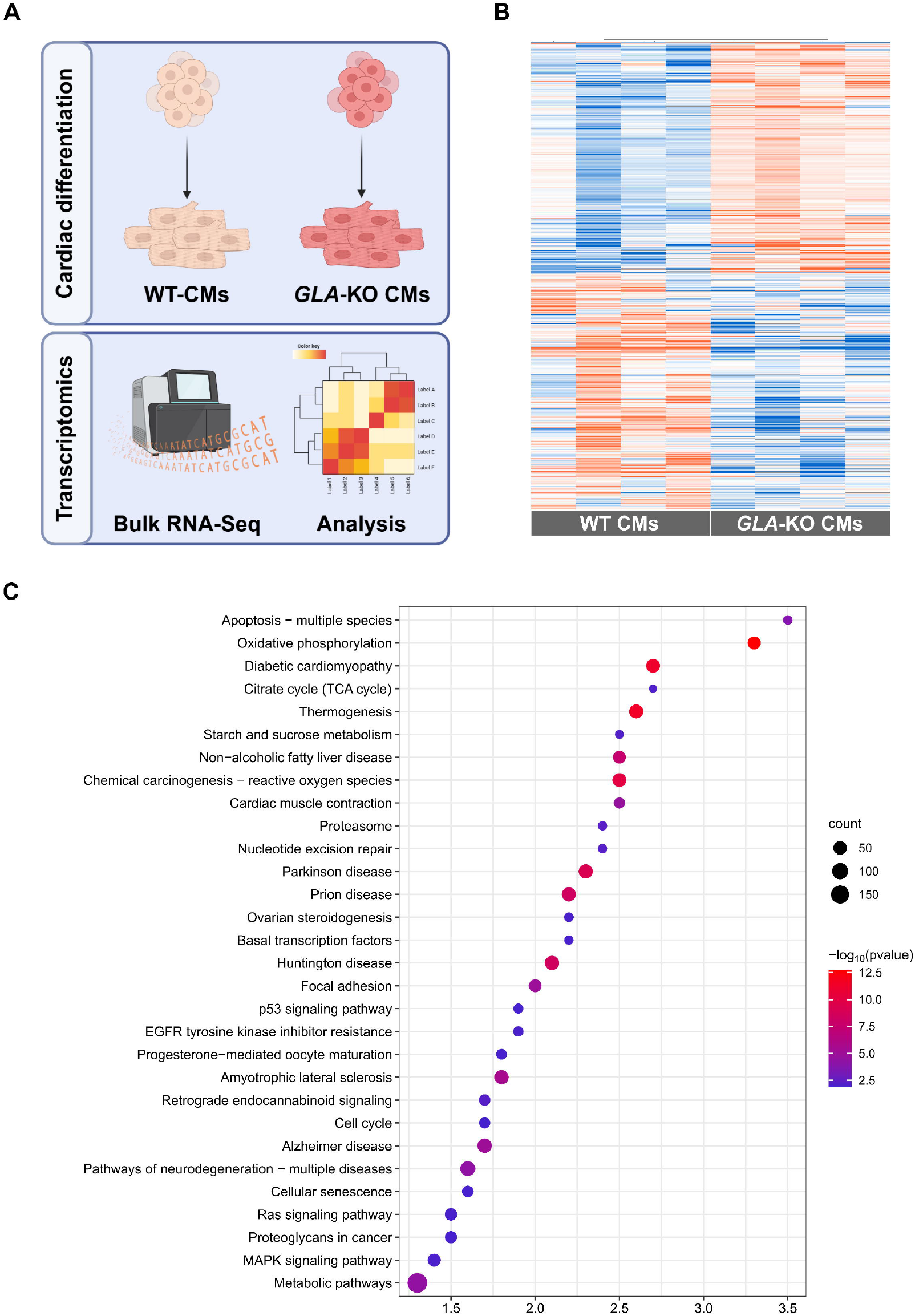
Comparative transcriptomics mirrors key Fabry disease phenotypes. (A) Schematic representation of the transcriptomics approach comparing day-65-old WT vs. *GLA*-KO CMs. Created in BioRender. Hoepfner, J. (2025) https://BioRender.com/85yv3f9. (B) A heatmap depicting the 2397 differentially expressed transcripts; (p ≤ 0.05 and base mean ≥ 5, n = 4 independent differentiations per group). (C) Bubble plot depicting the top 30 enriched KEGG pathways.

In essence, our transcriptomic approach both reflects well-established cellular phenotypes of Fabry disease and may provide new avenues of research.

### Nucleoside-modified α-galactosidase A mRNA therapy effectively restores α-galactosidase A enzyme activity in *GLA*-KO CMs

The available treatment options insufficiently manage Fabry disease, creating reasons to develop novel therapeutic strategies. A particularly promising modality that is steadily gaining pace and has generated first promising results in preclinical studies *in vitro*^[18]^ and *in vivo* both in mice and non-human primates^[17]^ is the use of *GLA* mRNA. However, as of now it remains unclear, whether *GLA* mRNA holds the potential to effectively address the archetypical cellular Fabry disease phenotypes, as the available animal models poorly reflect them, and *in vitro* studies are limited.

To address this question, we synthesized modGLA as a potential therapeutic intervention. The RNA was transcribed *in vitro* from a linearized plasmid template encoding the *GLA* open reading frame flanked by the alpha globin 5’-untranslated and beta globin 3’-untranslated region (Fig. 3 A). All uridines in the modRNA were substituted for with the modified nucleoside N1-methyl-pseudouridine (m1*Ψ*), and co-transcriptional capping was performed utilizing the Cap 1 analogue CleanCap AG. These modifications are known to reduce immunogenicity, while also allowing an enhanced translation efficiency^[26,27]^.

**Figure 3.**
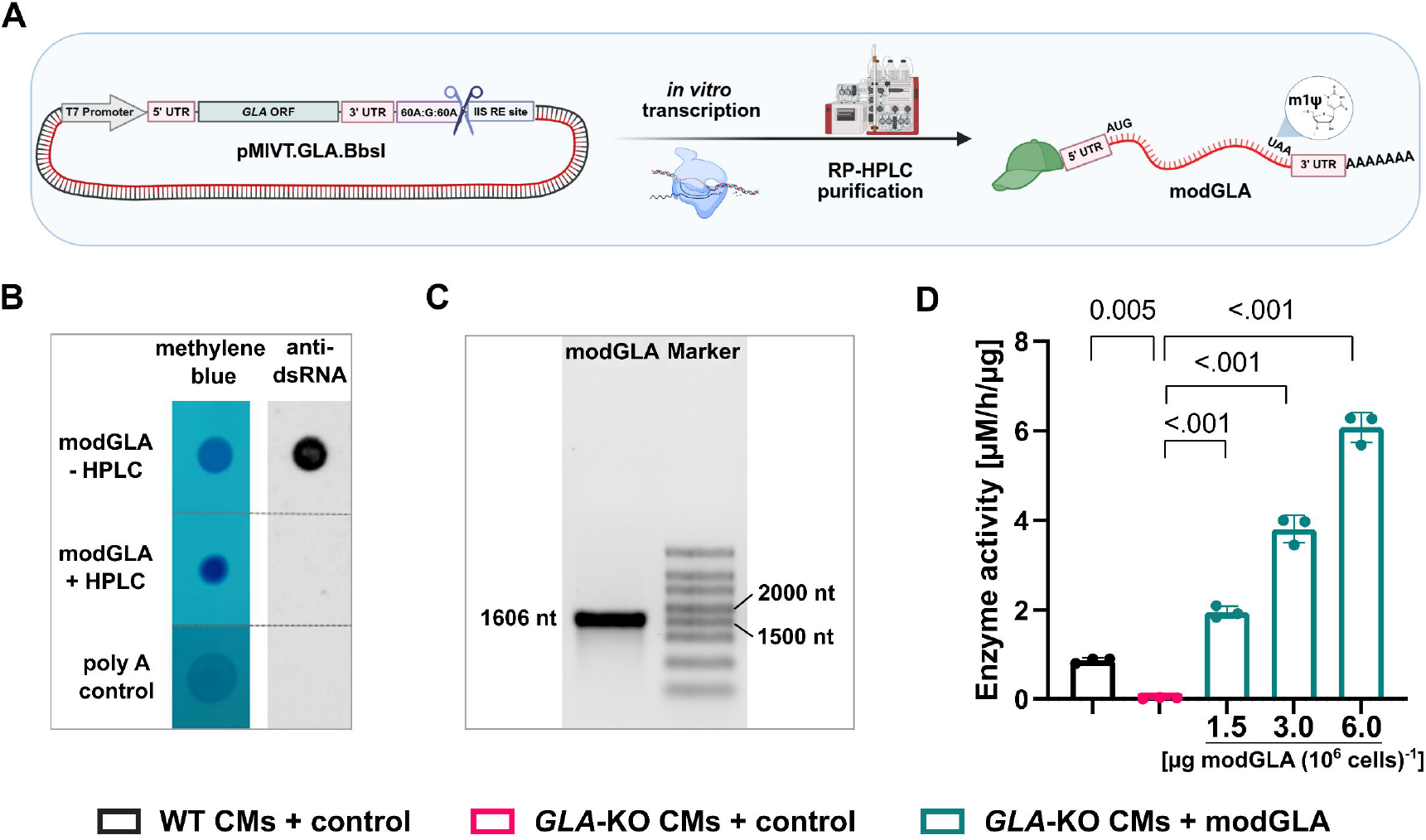
Nucleoside-modified *GLA* mRNA effectively restores α-galactosidase A enzyme function. (A) Production pipeline for nucleoside-modified *GLA* mRNA (modGLA) by in vitro transcription (IVT) from linearized plasmid templates followed by IP-RP-HPLC-based purification. Created in BioRender. Hoepfner, J. (2025) https://BioRender.com/wqdk2xg. (B) Dot Blot validating the successful removal of dsRNA byproducts from modGLA IVT products (modGLA - HPLC) by IP-RP-HPLC (modGLA + HPLC). (C) Analytical sodium hypochlorite agarose gel for analysis of purified modGLA. (D) Dose-dependent rescue of α-galactosidase A enzyme activity in *GLA*-KO CMs transfected with modGLA; (n = 3 independent differentiations, mean (SD), ordinary one-way ANOVA followed by Tukey’s multiple comparisons test, α = 0.05).

Subsequently, modGLA was purified using ion-pair reversed-phase high performance liquid chromatography (IP-RP-HPLC) (Fig. 3 A) which effectively eliminates dsRNA byproducts^[28]^ commonly found in T7 RNA polymerase transcription products^[29]^ (Fig. 3 B). Following RNA purification, denaturing sodium hypochlorite gel electrophoresis confirmed modGLA’s expected size of 1606 nucleotides, and its integrity (Fig. 3 C). Finally, transfecting increasing doses of modGLA into *GLA*-KO CMs dose-dependently rescued α-galactosidase A enzyme activity (Fig. 3 D). In the subsequent experiments, 6 µg modGLA per one million cells were applied since this dose produced the highest increase of α-galactosidase A enzyme activity without affecting cell viability, as measured by WST assay (Fig. S1).

### A-galactosidase A enzyme deficiency results in a cellular Fabry disease phenotype that is significantly rescued by modGLA therapy

While the molecular intricacies leading to the multifaceted cardiac manifestation of Fabry disease remain elusive, the trigger appears to be the massive deposition of α-galactosidase A glycosphingolipid substrates, including Gb3, within patients’ cells^[30]^. To investigate whether functional rescue of the α-galactosidase A enzyme activity and reduction of substrate levels is concomitant with a phenotypical rescue, we utilized modGLA as a therapeutic intervention. Following the administration of four weekly doses of therapeutic modGLA, or control modRNA (modfLuc, modEGFP, vehicle), the CMs were subjected to functional analyses at approximately 65 (± 5) days of age (Fig. 4 A). At this stage, we focused on well-established cellular phenotypes of Fabry disease that were also reflected in the earlier transcriptome analysis such as apoptosis, oxidative stress, and mitochondrial dysfunction. Evidently, modGLA therapy significantly rescued α-galactosidase A enzyme deficiency in *GLA*-KO CMs (Fig. 4 C). Automated fluorescence microscopic analysis of Gb3-associated signal revealed that this restored enzyme function significantly reduced Gb3 deposition within these cells, decreasing by on average 66.4% compared to modfLuc control-treated *GLA*-KO CMs (Fig. 4 B, D). In a second approach, we applied only a single dose of modGLA or modEGFP to 58-day-old CMs and again modGLA significantly rescued altered glycosphingolipid levels, specifically Gb3, galactobiosylceramide and nLc6, as analyzed by multiplex capillary gel electrophoresis coupled to laser-induced fluorescence detection; one week following treatment (xCGE-LIF; see Fig. S2).

**Figure 4.**
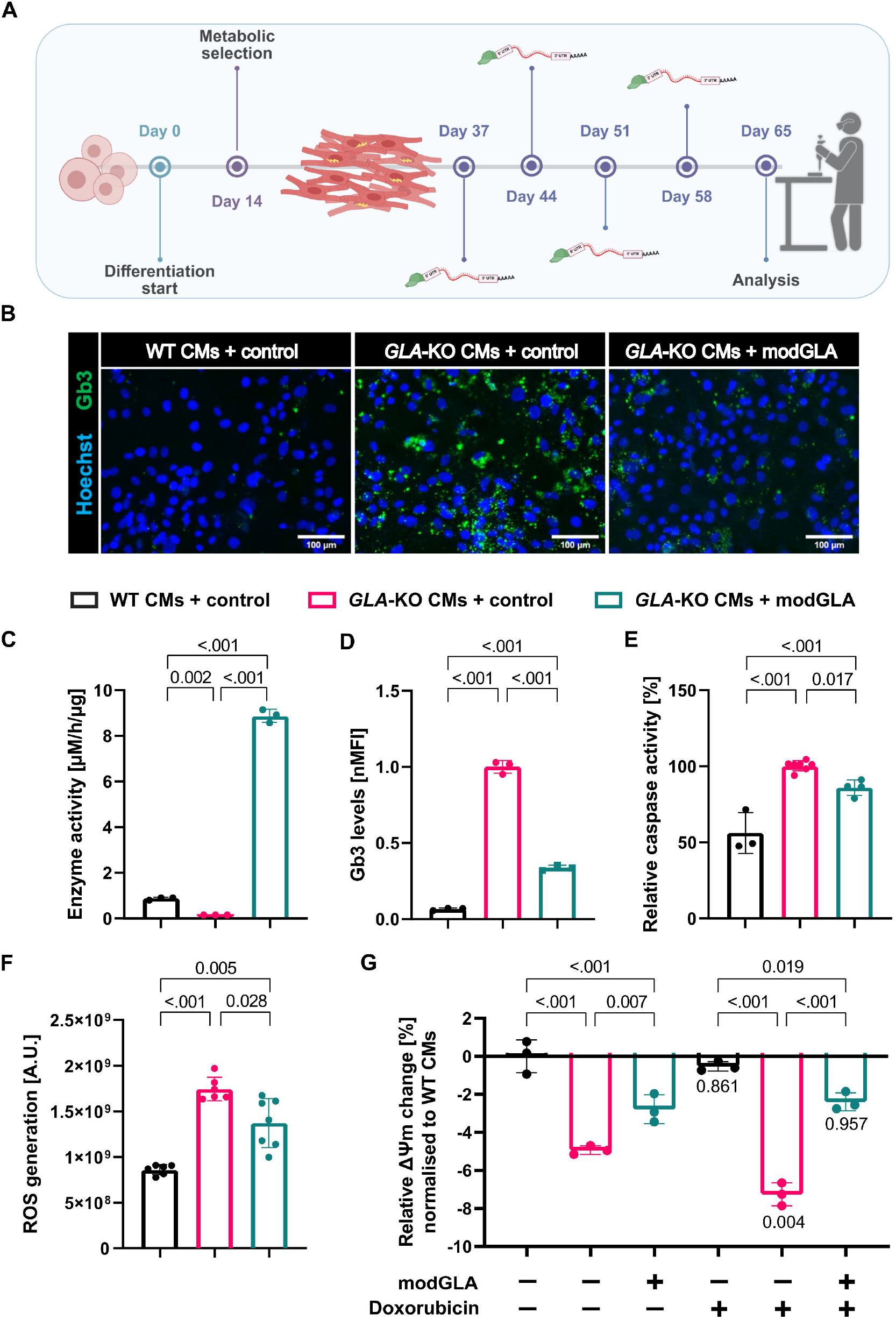
Glycosphingolipid accumulation and cellular disease phenotypes are rescued by modGLA therapy in Fabry cardiomyocytes. (A) Weekly treatment regimen of WT and *GLA*-KO CMs with modGLA, or control modRNA (modfLuc; modEGFP) or vehicle (only E). Time-points represent ranges of ± 5 days, while the intervals are consistently 7 days. The analyses in (B – G) were performed one week after the last modGLA dose. Created in BioRender. Bär, C. (2025) https://BioRender.com/x5t35jn. (B) Rescue of Gb3 accumulation in *GLA*-KO CMs by modGLA treatment as analyzed by fluorescence microscopy. Nuclei are stained by Hoechst33342. Scale bars represent 100 µm. (C) The α-galactosidase A enzyme activity as measured by α-galactosidase A enzyme activity assay in CM lysates; (n = 3 independent differentiations, mean (SD), ordinary one-way ANOVA followed by Tukey’s multiple comparisons test, α = 0.05). (D) Quantification of Gb3 signal in CMs from fluorescence microscopic images as those depicted in B, normalized to the number of nuclei, and expressed as “normalized mean fluorescence intensity (nMFI); (n = 3 independent differentiations, mean (SD), ordinary one-way ANOVA followed by Tukey’s multiple comparisons test, α = 0.05). (E) Relative caspase 3/7 activity measured by caspase-Glo® 3/7 assay in CMs challenged for 18 h with 2.56 µM doxorubicin; (n = 3-6 independent differentiations, mean (SD), ordinary one-way ANOVA followed by Tukey’s multiple comparisons test, α = 0.05). (F) Generation of reactive oxygen species (ROS) over a period of 6 h in CMs challenged with 100 µM H_2_O_2_ as measured by DCFDA assay in arbitrary units (A.U.); (n = 6-7 independent differentiations, mean (SD), Brown-Forsythe and Welch’s ANOVA followed by Dunnett’s T3 multiple comparisons test, α = 0.05). (G) Mitochondrial membrane potential (Δ*Ψ*m) relatively quantified in CMs with or without prior challenge with 1 µM doxorubicin for 24 h; (n = 3 independent differentiations, mean (SD), two-way ANOVA followed by Tukey’s multiple comparisons test, p-values below bars represent comparisons between means of equal color code, α = 0.05).

To assess the sensitivity of CMs to apoptotic stimuli, the cells were challenged with the cardiotoxic anthracycline doxorubicin prior to analysis. Strikingly, the changes observed in Gb3 levels were reflected in caspase 3/7 activities. Specifically, in comparison to WT CMs, control-treated diseased CMs exhibited significantly higher relative caspase 3/7 activity in response to the stress stimulus, which normalized significantly by approximately 32% in modGLA-treated *GLA*-KO CMs (Fig. 4 E). Additionally, we investigated the antioxidative capacity of our Fabry disease model by measuring the production of reactive oxygen species (ROS) in response to oxidative stress in form of hydrogen peroxide via H2DCFDA assay. As with the elevated caspase activity, control *GLA*-KO CMs also exhibited the greatest increase in ROS levels, which again normalized significantly by on average 41.8% upon modGLA treatment (Fig. 4 F). Finally, we used the fluorescent dye tetramethyl rhodamine ethyl ester (TMRE) in conjunction with flow cytometry to indirectly measure the mitochondrial membrane potential (Δ*Ψ*m) of CMs (Fig. 4 G). Interestingly, already in unchallenged CMs we observed a significant depolarizing shift in the Δ*Ψ*m in control *GLA*-KO CMs relative to control WT CMs that was partly rescued by modGLA treatment. However, after the cells were pre-treated for 24 hours with 1 µM doxorubicin, we observed this depolarizing shift to worsen in the control *GLA*-KO CMs. Such change was not observed in the control WT CMs or the modGLA-treated diseased CMs.

In summary, Fabry-archetypical phenotypes were confirmed at a functional level in our *in vitro* model, and for the first time, we demonstrated their rescue upon modGLA treatment.

### *GLA*-KO CMs display accelerated decay parameters of calcium transients and increased phospholamban phosphorylation

Pathway enrichment analysis performed with the significantly DEGs between *GLA*-KO and WT CMs, identified “cardiac muscle contraction” as a potentially relevant KEGG pathway. Because myocardial contraction is largely controlled by the spatio-temporal regulation of calcium ion concentrations^[31]^, we performed calcium imaging using the ratio metric calcium sensor Fura2-AM. The analysis of recorded calcium transients revealed a significantly increased decay velocity concurrent with a decreased 50% decay time in the diseased CMs, which normalized upon modGLA treatment (Fig. 5 A, B). A similar trend was observed with 7.9% of *GLA*-KO CMs displaying arrhythmic calcium transients, dropping to only 1.7% of cells following modGLA therapy (Fig. 5 C, S3). The calcium decay parameters are indicative of the reduction phase of temporarily higher cytoplasmic calcium concentrations during cardiac excitation-contraction coupling (Fig. 5 D). This led us to hypothesize a gain-of-function in the mechanisms shuttling calcium ions either out of the cell (NCX1) or back into the sarcoplasmic reticulum (via SERCA2a, regulated by its reversible inhibitor PLN). Strikingly, we observed no significant changes in the protein levels of NCX1 or SERCA2a between the disease model and respective controls (Fig. 5 E, F, G). However, the relative levels of PLN phosphorylated by cAMP-dependent protein kinase A (PKA) at serine 16 (pSer16-PLN), were significantly higher in the diseased CMs (Fig. 2E, H). This post-translational modification is known to relieve PLN’s inhibitory effect on SERCA2^[32]^. Notably, the hyperphosphorylation of PLN normalized in response to modGLA therapy.

**Figure 5.**
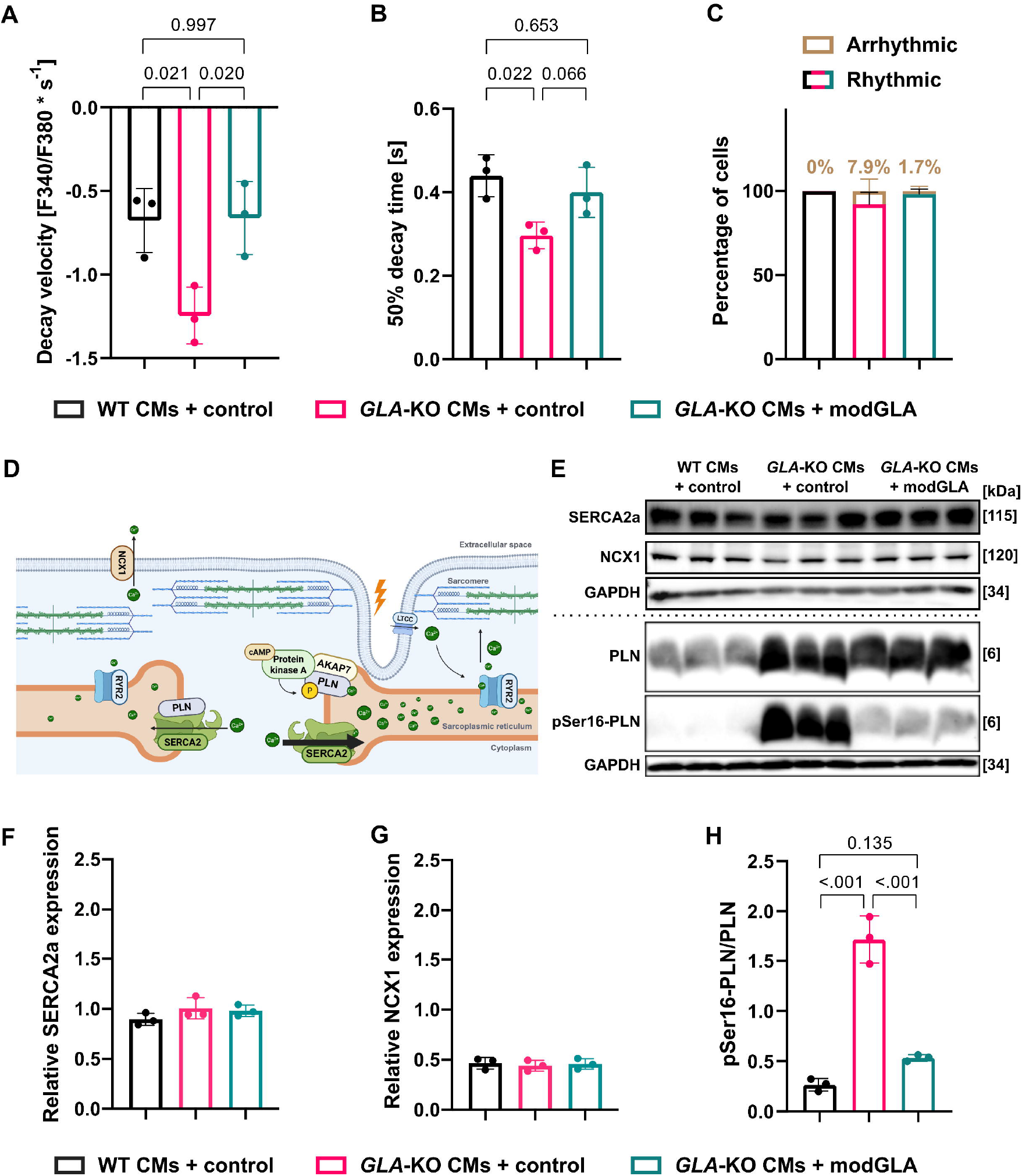
Enhanced calcium handling and hyperphosphorylated PLN in Fabry cardiomyocytes is partly normalized by modGLA therapy. Calcium imaging in CMs performed using the ratio metric calcium indicator Fura 2 to determine the (A) decay velocity of calcium transients as analyzed by the change in the ratio metric Fura 2 fluorescence in seconds [F340/380*s^-1^] and (B) the 50% decay time [s]; (n = 3 independent differentiations, calcium transients of 18-39 cells per replicate and group were recorded, mean (SD), ordinary one-way ANOVA followed by Tukey’s multiple comparisons test, α = 0.05; performed on log-transformed data after adding an offset constant [2] if values were previously negative). (C) Fraction of cells [%] showing arrhythmic calcium transients during calcium imaging; (n = 3 independent differentiations, calcium transients of 18-39 cells per replicate and group were recorded). (D) Schematic depiction of cardiac excitation-contraction coupling and the fine-tuning of calcium dynamics by the reversible SERCA2a-inhibitor phospholamban (PLN). Created in BioRender. Bär, C. (2025) https://BioRender.com/dc4kxkr. (E) Protein levels of SERCA2a, NCX1, total PLN, and PLN phosphorylated at serine 16 (pSer16-PLN) relative to GAPDH, analyzed by Western blot. The dotted line indicates two individual membranes. After detecting SERCA2a or pSer16-PLN, the respective membrane was stripped and reprobed for NCX1 or PLN, respectively. (F - H). Densitometric quantification of (F) SERCA2a, (G) NCX1, as well as (H) pSer16-PLN over total PLN; relative to GAPDH detected on the corresponding membranes depicted in (E); (n = 3 independent differentiations, ordinary one-way ANOVA followed by Tukey’s multiple comparisons test; α = 0.05).

Furthermore, we observed decreased trends of 90% decay time, decay rate *τ* and diastolic calcium in the Fabry CMs (Fig. S3 A, B, D). In contrast, the maximum rise velocity and calcium amplitude appeared to be increased in the disease model (see Fig. S3 C, E). Interestingly, the described alterations normalized upon modGLA treatment.

### Comparative transcriptomics in Fabry patient-derived CMs identifies cAMP and calcium signaling as drivers of cardiac manifestations

To validate the key findings observed in our generalized, patient-independent Fabry disease model in a disease-relevant, patient-specific context, we next employed an additional hiPSC-based *in vitro* model (F01 hiPSC), derived from a male Fabry patient carrying the c.959A > T missense mutation in *GLA*^[33]^. The corresponding isogenic control hiPSC line (F01-IC) was generated by prime editing CRISPR/Cas technology. Specifically, we differentiated all hiPSCs into CMs as described before and subjected both, the patient-derived (F01 vs. F01-IC CMs) and patient-independent CMs (*GLA*-KO vs. WT CMs), to bulk-RNA sequencing to compare the transcriptional profile of the two models (Fig. 6 A). This approach allowed us to assess, which molecular changes could be Fabry-distinctive, by focusing particularly on those DEGs with directionally similar expression changes (highlighted in green in Fig. 6 B; reciprocally regulated transcripts are marked red). Strikingly, when performing enrichment analysis on this subset of DEGs, cAMP and calcium signaling were amongst the top 5 enriched KEGG pathways (Fig. 6 C). Additionally, in accordance with the initial pathway analysis (see Fig. 2 C), DEGs with similar regulation patterns between the two models were particularly enriched in Ras/MAPK signaling pathways and the term “proteoglycans in cancer”. On protein level, and consistent with our initial results in patient-independent *GLA*-KO CMs, we found PLN to be hyperphosphorylated at serine 16 in patient-derived F01 CMs. Strikingly, this observation was partially reversed upon modGLA treatment (Fig. 6 D-G).

**Figure 6.**
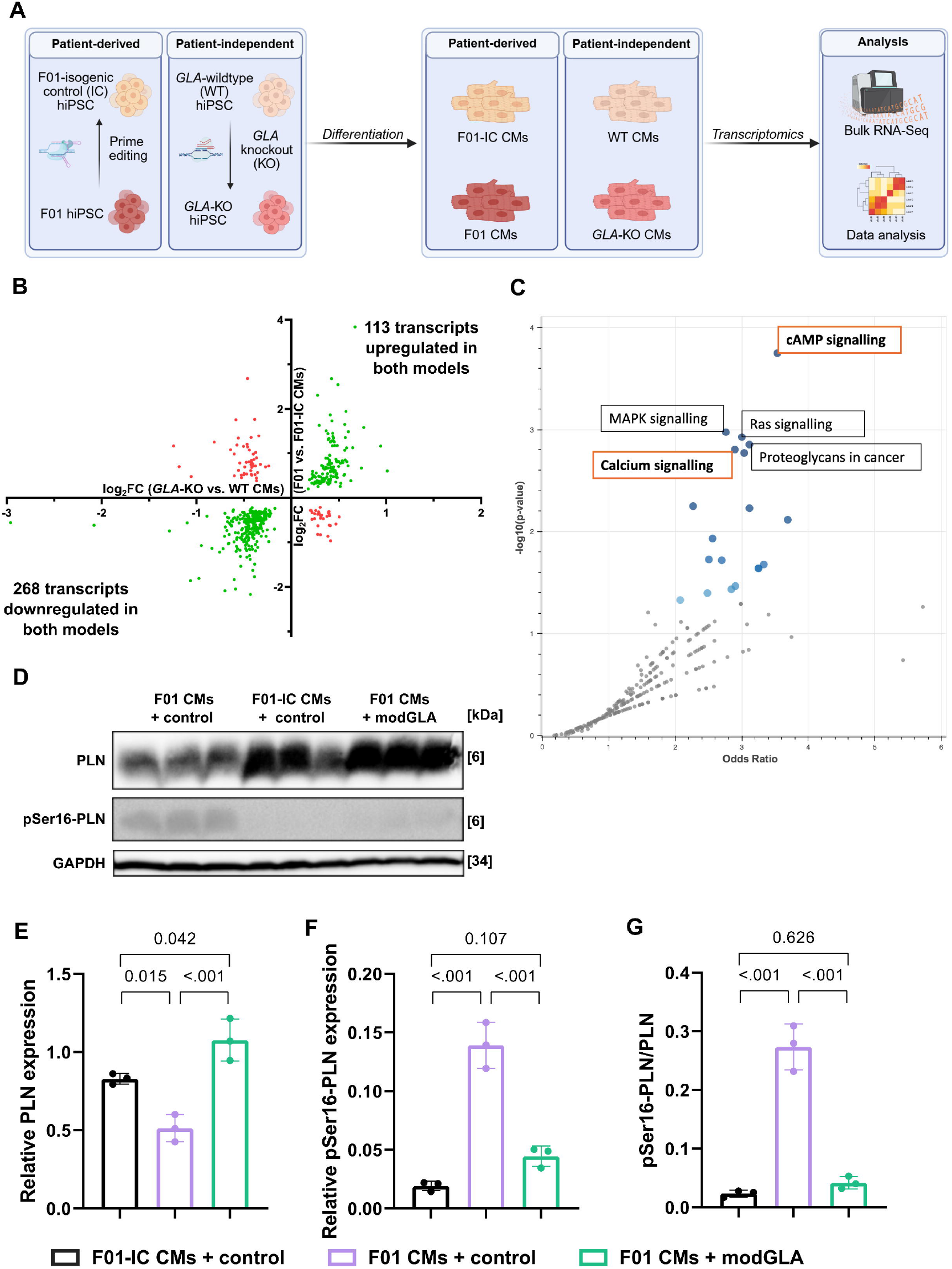
Comparative transcriptomics unravels Fabry-distinctive KEGG pathway signature. (A) Illustration of the combined approach applied to compare the transcriptional profile of two independent Fabry disease models, one patient-derived (F01 (n = 3 independent differentiations) vs. F01-IC CMs (n = 4 independent differentiations)), the other patient-independent (*GLA*-KO CMs vs WT CMs, n = 4 independent differentiations). Created in BioRender. Hoepfner, J. (2025) https://BioRender.com/85yv3f9. (B) Cartesian plot depicting the significantly differentially expressed genes (p ≤ 0.05) of two bulk-RNA sequencing data sets comparing F01 vs. F01-IC CMs (y-axis) and *GLA*-KO vs WT CMs (x-axis), respectively. Directionally similar regulated transcripts are highlighted in green, those transcripts reciprocally regulated between the disease models are labelled red. (C) Volcano plot displaying KEGG pathways enriched for the 381 transcripts showing directionally similar regulation in both disease models. Significantly enriched pathways are highlighted in blue. (D) Total PLN and pSer16-PLN protein levels as analyzed by Western Blot relative to GAPDH, detected on the same membrane. (E-G) Densitometric quantification of the protein levels detected in (D) relative to GAPDH; (n = 3 independent differentiations, ordinary one-way ANOVA followed by Tukey’s multiple comparisons test; α = 0.05).

In conclusion, by employing an independent patient-derived model of Fabry cardiomyopathy, we could validate the newly identified altered cAMP signaling and calcium handling, evidenced by PLN hyperphosphorylation, as a Fabry distinctive phenotype.

## Discussion

Here we report that loss of *GLA* entails several cellular disease phenotypes in hiPSC-derived CMs previously associated with Fabry disease, highlighting the validity of our disease model. Specifically, we observed an increased sensitivity to pro-apoptotic stimuli, an impaired capacity to neutralize reactive oxygen species, as well as an impaired mitochondrial membrane potential that deteriorated upon doxorubicin stress. In Fabry patients, glycosphingolipid build-up is proposed to activate secondary metabolic processes, which over time can manifest as Fabry cardiomyopathy^[30]^. The increased sensitivity of Fabry CMs to stress stimuli such as doxorubicin and H_2_O_2_ demonstrated in this study, suggests that the underlying cellular dysfunction in Fabry-affected cells might increase their susceptibility to deleterious factors encountered throughout each patient’s lifetime - particularly during periods of infection, inflammation, or the exposure to dietary or pharmacological toxins. Consequently, the ill-defined genotype-phenotype correlation in Fabry disease^[34]^, may in part be a result of non-genetic factors such as the unique lifestyle of each patient, which includes varying levels of these stressors. Future longitudinal and prospective cohort studies could aid in investigating this hypothesis. In addition, the great phenotypical heterogeneity of Fabry disease patients^[34]^, could potentially be deciphered by identifying putative confounding genetic variants by exome sequencing.

Enhanced calcium handling, in particular an increased amplitude and rising slope, as well as an elevated calcium load in the sarcoplasmic reticulum, had been reported before by Birket and colleagues in Fabry hiPSC-CMs^[35]^. Similarly, we observed enhanced calcium decay parameters in *GLA*-KO CMs. However, a mechanistic explanation for these observations was lacking thus far. Here we propose a first explanation for these calcium handling abnormalities in form of PLN hyperphosphorylated by PKA. Importantly, SERCA2a activity is tightly regulated by PLN, its reversible inhibitor. This inhibitory effect can be relieved by post-translational modifications, including phosphorylation at serin 16 by PKA^[32]^. Consequently, the consistently higher proportion of phosphorylated PLN over total PLN in these models, suggests a disinhibition of PLN’s inhibitory effect on SERCA2a function. In result, higher activity of SERCA2a in Fabry CMs likely entails a faster removal of calcium ions from the cytoplasm back into the sarcoplasmic reticulum during cardiac excitation-contraction coupling. This mechanism is in line with the increased calcium load in the sarcoplasmic reticulum observed by Birket and colleagues^[35]^. Congruently, we observed trends of lower diastolic calcium concentrations and higher calcium transient amplitudes in *GLA*-KO CMs (see Fig. S3 D, E). The upstream pathways leading to the phosphorylation of PLN’s Ser16 residue by PKA shall be a focus of future research efforts. Factoring in that the transcriptomics approach comparing the two independent disease models highlighted cAMP and Ras/MAPK signaling, two pathways that are known to collectively modulate cardiac calcium homeostasis (e.g.,^[36,37]^), investigations into their molecular interplay in the context of Fabry cardiomyopathy might prove particularly productive. The comparative pathway analysis also highlighted calcium signaling in general, substantiating an important role in the cardiac manifestation of Fabry disease. As to what degree this is the case remains to be answered by future studies.

Huurne et al. have recently demonstrated that *GLA* mRNA can restore α-galactosidase A enzyme activity in Fabry hiPSC-CMs and reduce Gb3 accumulation with an RNA dosing regimen every other day from 14 until 21 days post start of differentiation^[18]^. To further extend these findings, we utilized more mature hiPSC-CMs, as these cells are reported to develop substantial expression of key ionic currents and hence electrophysiological and calcium handling ability only after prolonged culture of >50 days^[38]^. Accordingly, we performed the functional analyses 65 days (± 5 days) after differentiation onset. To this date, it remained unknown whether modGLA therapy has the potential to clear Gb3 accumulation from CMs at that age. Notably, by successfully employing modGLA both as a weekly therapeutic intervention and a single dose treatment in relatively mature *GLA*-KO CMs, we provided first proof that glycosphingolipid deposition can be effectively reduced in CMs even after it manifested substantially. Since diagnosis in patients often comes late after glycosphingolipid accumulation and resulting symptoms already became evident^[39]^, this observation may be of clinical relevance. The effective weekly dosing regimen used in our study further corroborates the translational potential of this approach. However, whether modGLA proves more effective compared to ERT in eliminating substrate deposition from cardiomyocytes *in vivo* remains to be determined. Nevertheless, the consistent partial restoration of the investigated disease phenotypes in response to modGLA therapy confirms their causative relationship with α-galactosidase A enzyme deficiency and Fabry disease.

The partial rescue of these phenotypes might be explained by an incomplete removal of Gb3 deposition from the CMs. While longer treatment periods might clear the residual Gb3, it is conceivable that lysosomes packed with Gb3 become dysfunctional to a degree that might hinder their removal by the autophagic-lysosomal system or trafficking of functional α-galactosidase A into these compartments. Aptly, an impaired autophagic-lysosomal function has been described before by Song and colleagues in a model of Fabry cardiomyopathy^[40]^.

Expanding on the investigations in present study, future studies could increase the total treatment duration to elucidate whether glycosphingolipid deposits can be entirely cleared from affected cells, even when treatment is started after glycosphingolipid accumulation has manifested already. Conversely, more frequent dosing regimens may offer limited benefits, since systemic delivery of for instance lipid nanoparticle-encapsulated modGLA via intravenous injections several times a week, is not conducive to clinical translation.

Newer approaches such as engineered circular coding RNA^[41]^ might be explored instead to increase the half-life of the therapeutic RNA and hence reduce dosing frequency, which would not only limit costs but might additionally aid patient compliance.

However, it is conceivable that irreversible cellular damage may have manifested by the time treatment is initiated, which cannot be ameliorated by complete substrate removal alone. Such long lasting changes could for instance be epigenetic changes to patient DNA, as have been reported before^[42]^. In this case additional therapeutic strategies need to be developed that specifically target these altered pathways.

In summary, we provide mechanistic insight into the early cellular manifestation of Fabry cardiomyopathy, particularly in form of abnormal calcium handling resulting from PLN hyperphosphorylation. In addition, our data illustrate that mRNA therapy can rescue archetypical cellular disease phenotypes in Fabry hiPSC-CMs, supporting its potential as a future therapeutic strategy for Fabry disease.

## Material and methods

### Maintenance and differentiation of hiPSCs

HiPSCs were maintained in E8 medium (as described in Chen et al.^[43]^: DMEM F12 HEPES, 543 mg/l NaHCO_3_, 20 mg/l insulin, 10.7 mg/l holo-transferrin, 2 µg/l TGFβ1, 64 mg/l ascorbic acid 2-phosphate, 14 µg/l sodium selenite, 20 µg/l bFGF) and grown to 90-100% confluency on GelTrex-coated (25 µg/cm^2^) cell culture plates before passaging every 4-5 days into E8 medium supplemented with 2 µM Thiazovivin using 480 mM EDTA in DPBS. Differentiation into CMs was performed as described before^[44]^ by passaging the hiPSCs in clusters and subjecting them to biphasic Wnt pathway modulation (in RPMI1640 and GlutaMAX™ with HEPES, 0.5% (w/v) recombinant human albumin, 0.2% (w/v) L-ascorbic acid 2-phosphate sesquimagnesium salt hydrate: two days supplemented with CHIR99021, two consecutive days supplemented with IWP-2, and finally 4 days medium only). Eight days after start of differentiation, spontaneous contractions started and medium was switched to RPMI1640 and GlutaMAX™ with HEPES, 2% (w/v) serum-free B27 supplement for one week before performing metabolic selection in RPMI1640 without glucose, 0.5% (w/v) recombinant human albumin, 0.2% (w/v) L-ascorbic acid 2-phosphate sesquimagnesium salt hydrate, 4 mM for another week. Selected hiPSC-CMs were maintained in RPMI1640 and GlutaMAX™ with HEPES, 2% (w/v) serum-free B27 supplement.

### Alpha-galactosidase enzyme activity assay

The α-galactosidase A enzyme activity was determined as described before^[45]^, with few modifications. In brief, cell lysates were prepared for 30 minutes at 4°C on a rotating device using basal buffer (46 mM sodium phosphate dibasic, 27 mM sodium citrate, pH 4.6) with 0.5% Triton X-100. The lysates (20 µl) were incubated in black microtiter plates with 40 µl assay buffer (basal buffer with 117 mM N-acetyl-D-galactosamine and 6 mM 4-methylumbelliferyl-α-D-galactopyranoside) at 37°C for 1 hour. Next, the reaction was stopped by adding 70 µl of 400 mM glycine at pH 10.8. After adding a 4-methylumbelliferone standard (0.5 – 100 µM) to the plate, the fluorescence was detected using a Synergy HT plate reader (365/488 nm, Agilent Technologies). The α-galactosidase A enzyme activity was calculated from the standard curve and the known protein concentration of samples determined by Pierce BCA protein assay kit, as µM of 4-methylumbelliferone/h/µg of protein.

### Immunocytochemistry

Following a wash in DPBS, cultured cells were fixed in 4% PFA for 15 minutes at room temperature (RT). Permeabilization was performed for 20 minutes at RT (DPBS + 0.5% Tween-20, 0.1% Triton X-100, 0.1% Igepal CA-630), before staining the cells overnight at 4°C with the respective primary antibodies (Gb3: 1:500, #A2506, TCI; cTnT: 1:400, #ab45932, Abcam) diluted in DPBS + 1% BSA + 0.3% Triton X-100. The following day, staining with the secondary antibodies (Anti-mouse IgG Alexa Fluor 488: 1:500, #A-21202; Anti-rabbit IgG Alexa Fluor 594: 1:500, #A-21207; Thermo Fisher Scientific) and Hoechst33342 (1:2,000) was performed for 1 hour at RT. Microplates were imaged using a Cytation 1 device, while glass slides were first mounted in Vectashield HardSetTM Antifade Mounting Medium and then imaged on a Keyence BZ-X810 fluorescence microscope.

### Flow cytometry

To determine the differentiation efficiency of hiPSCs into derived CMs, flow cytometry was utilized. In brief, hiPSC-CMs were dissociated into single cells using 0.25X Trypsin/EDTA for 5 minutes at 37°C, then pelleted at 300 x g for 5 minutes and fixed in 4% PFA for 20 minutes at RT. Following two washing steps in DPBS, 10^6^ cells were permeabilized (DPBS + 1% BSA + 0.1% Triton X-100) per sample. Staining was performed for 1 hour at RT with either isotype control (REA Control Antibody, human IgG1, FITC, REAfinity; #130-113-449, Miltenyi) or anti cardiac troponin T antibody (Cardiac Troponin T Antibody, anti-human/mouse/rat, FITC, REAfinity; #130-119-674, Miltenyi) diluted 1:50 in DPBS + 1% BSA + 0.1% Triton X-100. Following two washes, cells were pelleted and resuspended in DPBS + 1% BSA + 2 mM EDTA to approximately 10^6^ cells/ml. Analysis was performed using a CytoFLEX S flow cytometer (Ex/Em: 488/517 nm).

### Transmission electron microscopy

Cells were grown on membranes in well inserts (Greiner Bio-One 657651, 0.4 µm pore size thin certs) and fixed by submersion in 150 mM HEPES buffer at pH 7.35 containing 1.5% formaldehyde and 1.5% glutaraldehyde at RT for 30 minutes and then over night at 4°C. The membrane was cut off the insert and postfixed and stained in 1 % osmium tetroxide for 2 h at RT and then over night at 4°C in 1 % uranyl acetate, both aqueous solutions. After dehydration in acetone, samples were embedded in EPON and polymerized between aclar foils. The foil was removed afterwards, and the samples glued on EPON blocks and cut parallel to the cells. 50 nm sections were poststained with uranyl acetate and lead citrate^[46]^ and observed in a Zeiss EM 900 equipped with a side mounted camera (TRS, Moorenweis).

### Transcriptome analysis

To perform bulk-RNA sequencing, total RNA was extracted using the miRNeasy mini kit (Qiagen) from hiPSC-CMs according to the manufacturer’s instructions for RNA isolation from cultured cells. To validate a comparable RNA quality of samples, the RNA integrity was determined using a Nano 6000 kit on a Bioanalyzer 2100 (Agilent Technologies). The RNA sequencing and preceding directional library preparation was performed by NovoGene Co. In brief, total RNA (500 ng per sample) was rRNA depleted and then isolated using ethanol precipitation. Following fragmentation, first strand cDNA synthesis was performed using random hexamer primers. Second strand cDNA synthesis was performed while substituting dUTPs for dTTPs. Next, A-tailing was utilized and after that adapter ligation, size selection, USER enzyme digestion and amplification followed by purification to finalize the library. Then, quality control of the library was performed using a Qubit 2.0 (Thermo Fisher Scientific), a Bioanalyzer 2100 and real-time PCR. Using an Illumina Novaseq 6000 device in conjunction with an S4 flow cell and v1.5 reagent, the pooled libraries were sequenced with a paired-end strategy (PE150). The processing and analysis of raw read data was performed using the galaxy.eu platform (v2.0.1). Specifically, *TrimGalore!* was used for raw read data quality trimming before mapping reads to the human reference genome GRCh38.p13; release 34 (GENCODE.org) via *RNAstar* or *Bowtie2*. Next, *feature counts* was used to derive raw read counts. Finally, *DeSeq2* (v1.42.0) was used with default settings for differential gene expression analysis. Finally, pathway analysis was performed using *DAVID*^[22]^ (David.abcc.ncifcrf.gov) or *Enrichr*^[47]^ (maayanlab.cloud).

### Gene correction of Fabry patient-derived human iPSCs using prime editing

*GLA* gene correction of the human iPSC line “MHHi029-A^[33]^” carrying the c.959A>T mutation, was performed using prime editing, based on transient transfection of the PB-PE plasmid system using Lipofectamine Stem (Thermo Fisher Scientific), as described elsewhere^[48]^. Specifically, pegRNA oligonucleotides were annealed and cloned into the BbsI-HF/BsaI-HFv2-linearized PB-PE backbone. The pegRNA consisted of the scaffold sequence (see Eggenschwiler et al.^[48]^), the reverse transcriptase template (CATCA<u>A</u>TCAGGACCC<u>G</u>TTGG; changes compared to target sequence underlined), and the pegRNA primer binding site (GCAAGCAAGGGTA).

### Production of nucleoside-modified mRNA (modRNA)

Nucleoside-modified mRNA encoding for firefly luciferase (modfLuc), human α-galactosidase A (modGLA) or enhanced green fluorescent protein (modEGFP) were produced by in vitro transcription. First, specific plasmid templates were generated from pMA-RQ backbones each encoding: CleanCap AG-compatible T7-promoter - human α-globin 5’ UTR – KOZAK – ORF of the gene of interest - human β-globin 3’ UTR – segmented poly adenine stretch (60A:G:60A; see Trepotec et al.^[49]^) – type II S restriction enzyme site (BspQI/BbsI/BtgzI). These plasmid templates (see supplemental information for sequence information) were linearized by restriction digestion and purified by general phenol/chloroform extraction before being transcribed in vitro into modRNA (for a 1 ml *in vitro* transcription reaction: 50 µg linearized pDNA, 8000 U T7 RNA polymerase, 1000 U RNase inhibitor, 2 U inorganic pyrophosphatase, 4 mM CleanCap AG, 5 mM each of ATP, CTP, GTP and m1*Ψ*, 4 mM Tris-HCl pH 8, 10 mM DTT, 2 mM spermidine, 0.002% Triton X-100, 16.5 mM magnesium acetate) at 37°C for 4-24 h. DNA templates were removed by TURBO DNase digestion (100 U/ml IVT reaction; 15 minutes at 37°C). The modRNA was ultrafiltrated with Amicon Ultra-4 centrifugal filters (10 kDa MWCO) before removing 5’-triphosphates by Antarctic phosphatase treatment (5 U/pmole of uncapped RNA ends assuming 95% capping efficiency; 30 minutes at 37°C). Purification was performed via IP-RP-HPLC on an Äkta pure 25 device in conjunction with a PLRP-S column at 62°C (Agilent Technologies: column volume (CV) = 4.16 ml; 4000 Å; 30 µm); or using the Monarch RNA Cleanup Kit (NEB, T2050, only applies to Fig. S2). After loading, the modRNA was washed with eluent A (100 mM TEAA in HPLC-grade H_2_O, pH 7) followed by a linear gradient of up to 20% eluent B (100 mM TEAA in HPLC-grade H_2_O, 25% (v/v) acetonitrile, pH 7). Elution was performed over 16.2 CVs with a flow rate of 2.8 ml/minute using a linear gradient from 20 to 61% eluent B. The final product was subjected to buffer exchange into 1 mM sodium citrate at pH 6.4 using Amicon Ultra-4 centrifugal filters (10 kDa MWCO) before validating RNA integrity and size on a sodium hypochlorite agarose gel^[50]^ and detecting dsRNA by Dot Blot as described before^[51]^; with the additions of cross-linking the RNA to the membrane by UV-light (3 minutes at 120,000 µJ/cm^2^) and validating equal loading by methylene blue staining (0.02% methylene blue in 0.3M sodium acetate).

### Transfection of modRNA

For transfection of modRNA into cultured cells, RNAiMAX (Thermo Fisher Scientific)-modRNA complexes were prepared following the manufacturer’s instructions in OptiMEM I medium, with 1 µg of RNA/µl of transfection reagent. 6 µg of modRNA were transfected per one million hiPSC-CMs. 5 hours after adding the transfection mix, it was replaced by full cell culture medium (RPMI1640 and GlutaMAX™ with HEPES, 2% (w/v) serum-free B27 supplement).

### Caspase activity assay

The relative caspase 3/7 activity was determined using the Caspase-Glo 3/7 kit (Promega) according to the manufacturer’s instructions. In brief, cells were seeded into Geltrex-coated microplates at 30,000 hiPSC-CMs /cm^2^ one week before performing the assay. Next, 18 hours preceding the assay, a doxorubicin (2.56 µM) stress stimulus, diluted in RPMI1640 and GlutaMAX™ with HEPES, 2% (w/v) serum-free B27 supplement, was applied to respective wells. To perform the assay, the caspase-Glo reagent was prepared and added to lysed cells. Following incubation for 1 hour at 37°C, luminescence was recorded with a Synergy HT plate reader (Agilent Technologies).

### Mitochondrial membrane potential measurement

The mitochondrial membrane potential was indirectly measured using the fluorescent dye tetramethyl rhodamine ethyl ester (TMRE). 24 hours prior to the assay, the respective cells were challenged with 1 µM doxorubicin in RPMI1640 and GlutaMAX™ with HEPES, 2% (w/v) serum-free B27 supplement. The hiPSC-CMs were dissociated into single cells (0.25% Trypsin/EDTA for 5 minutes at 37°C) and stained adhering to the manufacturer’s instructions provided with the TMRE-Mitochondrial Membrane Potential Assay Kit (Abcam). The cells were strained through a 100 µm strainer and analyzed on a CytoFLEX S (Beckman Coulter) cytometer (Ex/Em: 488/575 nm).

### Measurement of cellular ROS

Cellular ROS was measured via the DCFDA/H2DCFDA Cellular ROS Assay Kit (Abcam) following the manufacturer’s instructions. In brief, hiPSC-CMs were seeded onto GelTrex-coated microplates at 30,000 cells/cm^2^. At least one week later, staining was performed for 45 minutes at 37°C with 25 µM 2’,7’-dichlorofluorescin diacetate (DCFDA). Then, the cells were washed with DPBS before adding 100 µM H_2_O_2_ in assay buffer or buffer only as a background control. Finally, the plates were imaged every 30 minutes for 6 hours total using a Cytation 1 (Agilent Technologies) device (Ex/Em: 485/535 nm). Data were analyzed by subtracting the background from the raw read values and plotting the results to calculate the area under the curve.

### Calcium imaging

Calcium transients of individual Fura2-AM-loaded hiPSC-CMs, attached to MatriGel-coated 18 mm glass cover slips, were recorded on a dual excitation fluorescence photomultiplier setup (IonOptix). First, hiPSC-CMs were loaded with Fura2-AM (1.5 µM Fura2-AM in RPMI1640 medium containing GlutaMAX™ supplement with HEPES, 2% (v/v) serum-free B27 supplement) for 30 minutes at 37°C and 5% CO_2_. Following two 15-minute washing steps in medium only, cover slips were transferred to a custom-built perfusion chamber and paced at 0.5 Hz and 25 V (MyoPacer Cell stimulator; IonOptix) under constant flow of CM imaging solution (117 mM NaCl, 20 mM HEPES, 10 mM creatine, 10 mM D-glucose, 5.7 mM KCl, 5 mM sodium pyruvate, 1.25 mM CaCl_2_, 1.2 mM NaH_2_PO_4_, 0.66 mM MgSO_4_; pH 7.4 at 37°C). Calcium transients of randomly selected hiPSC-CMs that reacted to pacing, were recorded by detecting emitted fluorescence at 510 nm after excitation at both 340 and 380 nm. Finally, 20 calcium transients were averaged per cell using the IonWizard software (IonOptix, Version 6.5) and the parameters baseline ratio (R), ratio amplitude (R), maximum ratio increase velocity (R/s), time to peak (s), maximum decay velocity (R/s), 50 and 90% decay time (s) and exponential decay constant tau (s) were calculated. Outliers in the data were identified using ROUT (Q = 5%).

### Western Blot

Cells were lysed in RIPA buffer (+ protease & phosphatase inhibitor) by sonication. The protein concentration in lysates was determined by Pierce BCA protein assay kit (Thermo Fisher Scientific). Next, total protein (25 µg) was separated on an SDS polyacrylamide gel and then transferred onto a polyvinylidene membrane by wet blotting (Bio-Rad). The primary antibodies were used as follows to detect the respective proteins: GAPDH (1:20000, ab8245, Abcam); PLN (1:5000, A010-14, Badrilla); phospho-Ser16-PLN (1:5000, A010-12AP, Badrilla); SERCA2a (1:1000, MA3-919, Thermo Fisher Scientific); NCX1 (1:10000, ab177952, Abcam). Prior to reprobing, membranes were stripped in Restore PLUS Western Blot Stripping Buffer (Thermo Fisher Scientific).

### Statistics

Statistical analyses of data were performed using GraphPad Prism (v9.1.0). Data are presented as mean (SD). To compare means between two groups, an unpaired two-tailed Student’s t-test was used. For comparisons among three or more groups, ordinary one-way ANOVA was followed by Tukey’s multiple comparisons test when homogeneity of variances could be assumed. When variances were unequal, Brown-Forsythe and Welch’s ANOVA were used and followed by Dunnett’s T3 multiple comparisons test. For experiments involving two independent variables, two-way ANOVA followed by Tukey’s multiple comparisons test was applied. Post hoc p-values are only reported when the corresponding ANOVA indicated a statistically significant difference (p ≤ 0.05).

## Acknowledgements

We appreciate the outstanding technical support by A. Gietz from the Institute of Molecular and Translational Therapeutic strategies (IMTTS) at Hannover Medical School (MHH). We thank the Department of Bio- and Environmental Analytics at Fraunhofer Institute for Toxicology and Experimental Medicine (ITEM) for their excellent technical support and provision of HPLC infrastructure.

## Author contributions

MJ designed experiments, analyzed data, and drafted the manuscript. MJ, JHo, TT and CB conceptualized the study, and revised the manuscript. SE, LO, JLY contributed to the production of modRNA. NW provided significant support with calcium imaging. CJ, RE and TC planned and performed the prime editing-based gene correction of the patient-derived iPSC line. JHe performed electron microscopic analyses. KX performed bioinformatic analysis of transcriptomics data. JB and FB performed analysis of glycosphingolipids. All authors reviewed and approved the final version of the manuscript.

