## Supporting information for "Nucleoside-Modified mRNA Encoding Alpha-Galactosidase A Reverses Fabry Disease Phenotypes in Human IPSC-Derived Cardiomyocytes"

**This file includes:**

Supplemental methods

Ethics statement

Figures S1, S2 and S3

Nucleotide sequences of plasmids and transcribed modRNA

References

### Supplemental methods

#### WST-1 cell viability assay

The influence of modRNA transfection on cell viability of hiPSC-CMs was assessed by WST-1 assay (Roche, #5015944001) according to the manufacturer's recommendations.

#### Profiling of glycosphingolipid-derived glycans

Analysis of glycans derived from glycosphingolipids was performed by multiplexed capillary gel electrophoresis coupled to laser-induced fluorescence detection (xCGE-LIF) as described previously[1]. In brief, glycans were cleaved off glycosphingolipids using ceramide glycanase. The resulting free reducing ends were labelled with the fluorescent dye 8-aminopyrene-1,3,6-trisulfonic acid (APTS), which enabled glycan detection during the subsequent capillary gel electrophoresis. Glycan subtypes were identified by comparison of migration times to a migration time database containing the respective glycans.

### Ethics statement

All procedures involving human material were approved by the local ethics committee of Hannover Medical School (approval nos. 409 and 3668-2017). Human induced pluripotent stem cells (hiPSCs) in this study were obtained and used in accordance with institutional and national ethical guidelines. Donors provided informed consent for the generation and research use of hiPSC lines.

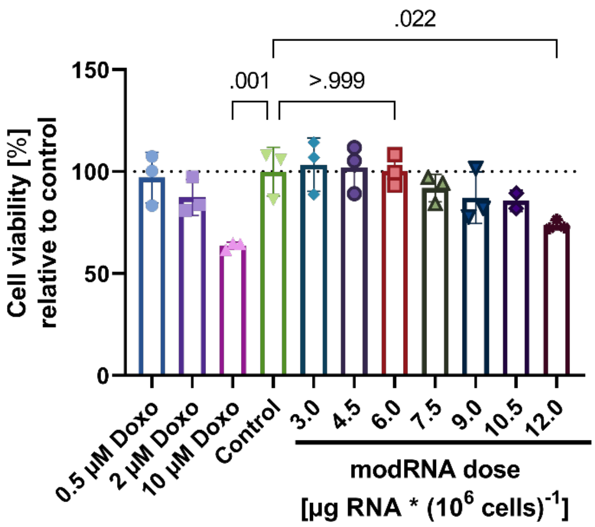


**Figure S1 – Cell viability of hiPSC-CMs in response to escalating doses of modGLA**

Cell viability of hiPSC-CMs as assessed by WST-1 assay 24 hours after transfection with increasing doses of modGLA or after doxorubicin treatment, which served as a positive control for cardiotoxic effects. The cell viability begins to drop off with modGLA doses of 7.5 µg RNA * ($10^6$ cells)$^{-1}$. Statistical analysis was performed using one-way ANOVA followed by Dunnett's multiple comparisons test, comparing the mean of each group with the control (mean (SD); n = 3 individual wells; α = 0.05).

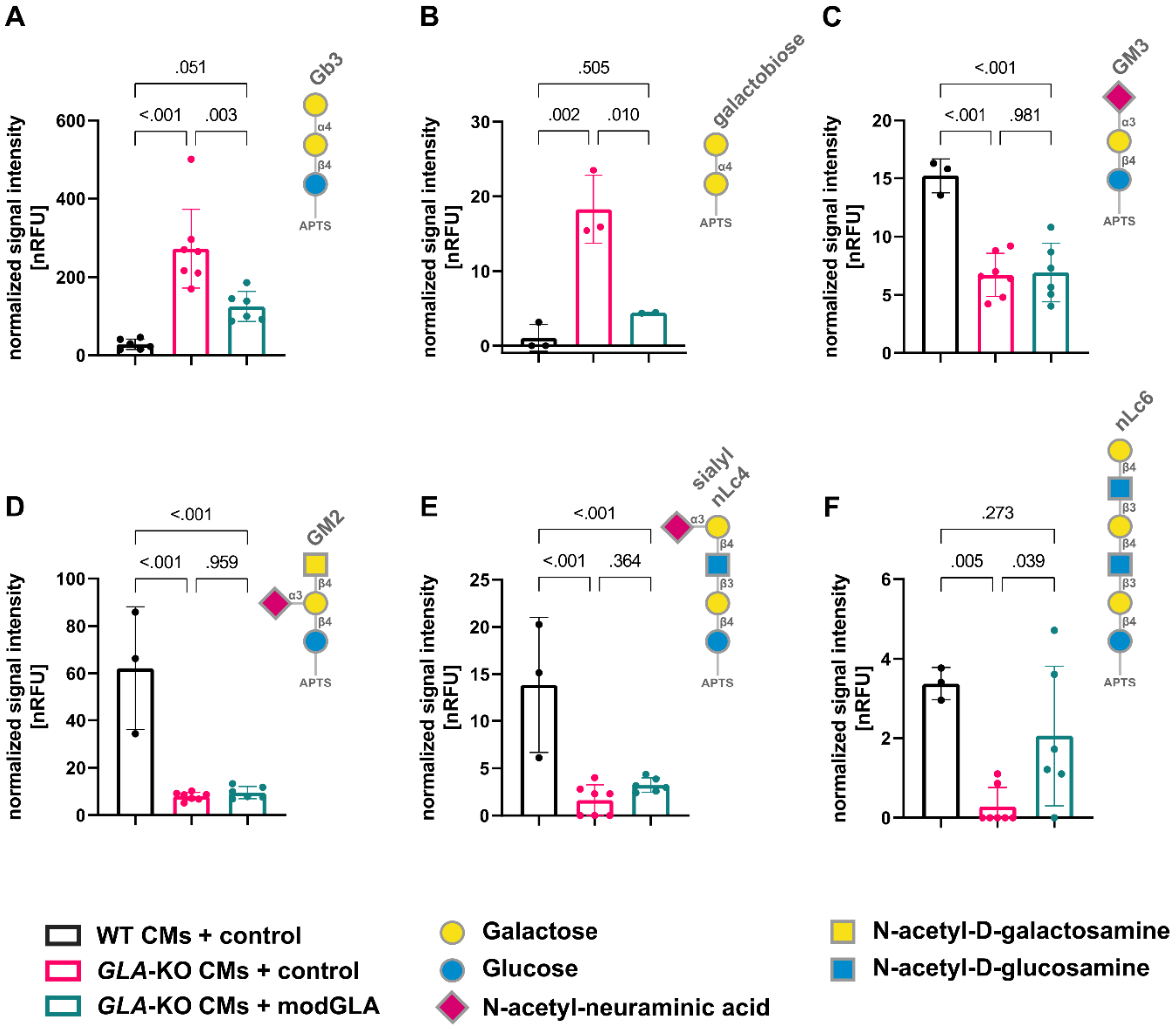


**Figure S2 - xCGE-LIF-based glycan profiling reveals altered glycosphingolipid composition in *GLA*-KO CMs**

Quantification of glycosphingolipid-derived glycans was conducted in 65-days-old WT and *GLA*-KO CMs (modEGFP control treated) and modGLA-treated *GLA*-KO CMs to assess the therapeutic effect of a single dose of modGLA; one week following transfection. Assigned glycans are presented as pictograms. A single dose of modGLA significantly normalized Gb3, galactobiose, and nLc6 glycan levels. (n = 2-7 independent differentiations, mean (SD), ordinary one-way ANOVA followed by Tukey's multiple comparisons test; $\alpha = 0.05$).

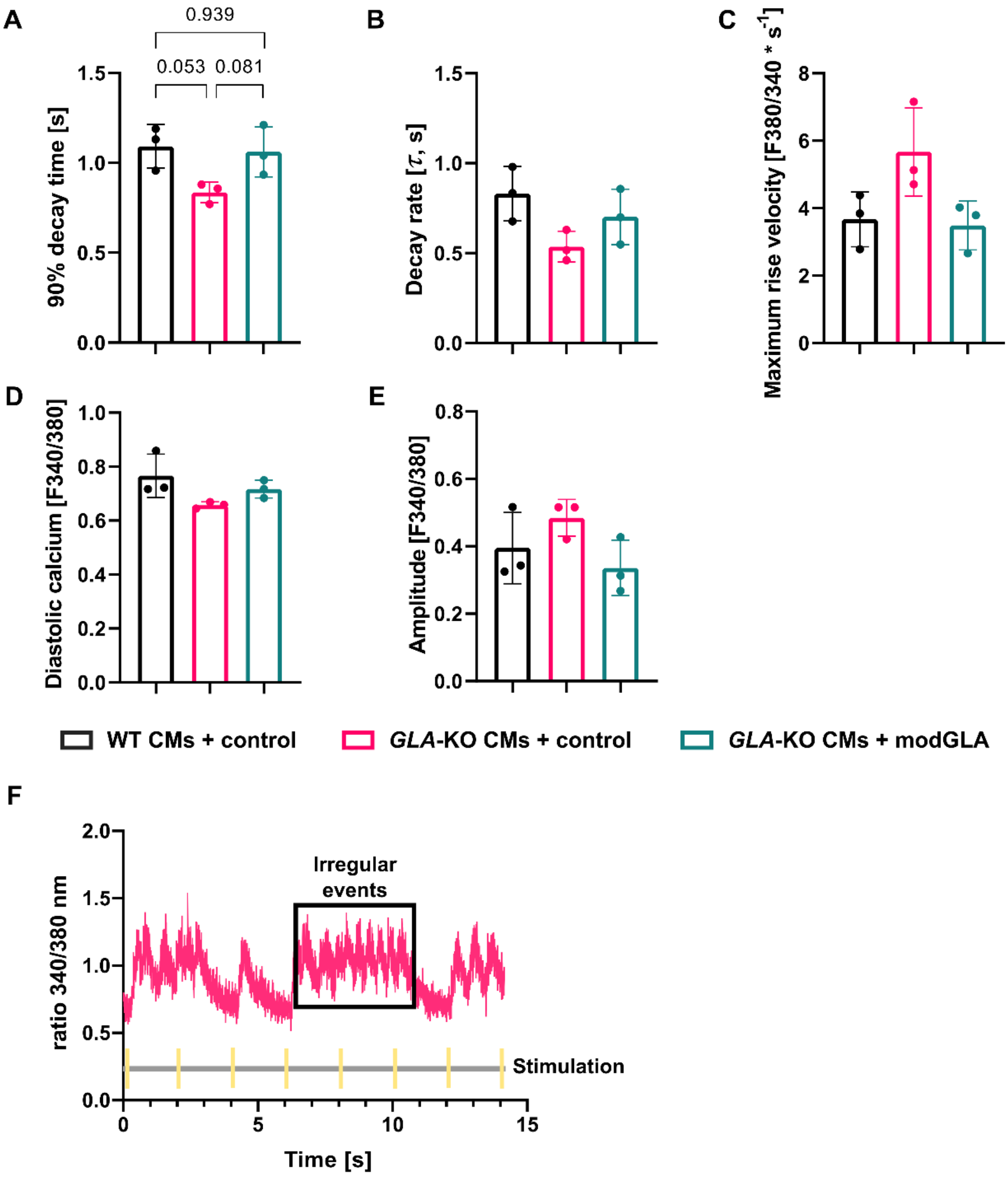


**Figure S3 – Potential signs of altered calcium handling in *GLA*-KO CMs**

Fura 2-based calcium imaging was used to determine the (A) 90% decay time of calcium transients [s], (B) the decay rate τ [s], (C) the maximum rise velocity as determined by the change in the ratio metric Fura 2 fluorescence per second [F340/380 $s^{-1}$], (D) the diastolic calcium levels and (E) the transient amplitude [F340/380], (n = 3 independent differentiations, calcium transients of 20-39 cells were recorded per replicate and group, mean (SD), ordinary one-way ANOVA followed by Tukey's multiple comparisons; $\alpha = 0.05$. Test performed on log-transformed data). (F) Exemplary depiction of arrhythmic calcium transients detected in a *GLA*-KO CM (highlighted by black box labelled "irregular events"). In contrast, a regular transient precedes the black box between electrical stimuli at 4 and 6 seconds.

**Nucleotide sequences of plasmids and transcribed modRNA**

U = N1-methyl-pseudouridine
Cap1 = (7-methyl-guanosine)-triphosphate linkage-(2-*O*-methyladenosine)

>pMIVT.GLA.BbsI

CTAAATTGTAAGCGTTAATATTTTGTTAAAATTCGGATGTGCTGCAAGGCGAGCGGCCGCAGGCCGTC
AAGGCCGCATGGTACCTGACAGCTGTAATACGACTCACTATAAGGAGAACTCTTCTGGTCCCCACAGA
CTCAGAGAGAACCCACCGGTGCCGCCACCATGCAGCTGAGGAACCCAGAACTACATCTGGGCTGCGCG
CTTGCGCTTCGCTTCCTGGCCCTCGTTTCCTGGGACATCCCTGGGGCTAGAGCACTGGACAATGGATT
GGCAAGGACGCCTACCATGGGCTGGCTGCACTGGGAGCGCTTCATGTGCAACCTTGACTGCCAGGAAG
AGCCAGATTCCTGCATCAGTGAGAAGCTCTTCATGGAGATGGCAGAGCTCATGGTCTCAGAAGGCTGG
AAGGATGCAGGTTATGAGTACCTCTGCATTGATGACTGTTGGATGGCTCCCCAAAGAGATTCAGAAGG
CAGACTTCAGGCAGACCCTCAGCGCTTTCCTCATGGGATTCGCCAGCTAGCTAATTATGTTCACAGCA
AAGGACTGAAGCTAGGGATTTATGCAGATGTTGGAAATAAAACCTGCGCAGGCTTCCCTGGGAGTTTT
GGATACTACGACATTGATGCCCAGACCTTTGCTGACTGGGGAGTAGATCTGCTAAAATTTGATGGTTG
TTACTGTGACAGTTTGGAAAATTTGGCAGATGGTTATAAGCACATGTCCTTGGCCCTGAATAGGACTG
GCAGAAGCATTGTGTACTCCTGTGAGTGGCCTCTTTATATGTGGCCCTTTCAAAAGCCCAATTATACA
GAAATCCGACAGTACTGCAATCACTGGCGAAATTTTGCTGACATTGATGATTCCTGGAAAAGTATAAA
GAGTATCTTGGACTGGACATCTTTTAACCAGGAGAGAATTGTTGATGTTGCTGGACCAGGGGGTTGGA
ATGACCCAGATATGTTAGTGATTGGCAACTTTGGCCTCAGCTGGAATCAGCAAGTAACTCAGATGGCC
CTCTGGGCTATCATGGCTGCTCCTTTATTCATGTCTAATGACCTCCGACACATCAGCCCTCAAGCCAA
AGCTCTCCTTCAGGATAAGGACGTAATTGCCATCAATCAGGACCCCTTGGGCAAGCAAGGGTACCAGC
TTAGACAGGGAGACAACTTTGAAGTGTGGGAACGACCTCTCTCAGGCTTAGCCTGGGCTGTAGCTATG
ATAAACCGGCAGGAGATTGGTGGACCTCGCTCTTATACCATCGCAGTTGCTTCCCTGGGTAAAGGAGT
GGCCTGTAATCCTGCCTGCTTCATCACACAGCTCCTCCCTGTGAAAAGGAAGCTAGGGTTCTATGAAT
GGACTTCAAGGTTAAGAAGTCACATAAATCCCACAGGCACTGTTTTGCTTCAGCTAGAAAATACAATG
CAGATGTCATTAAAAGACTTACTTTAAAGCTTGCTCGCTTTCTTGCTGTCCAATTTCTATTAAAGGTT
CCTTTGTTCCCTAAGTCCAACTACTAAACTGGGGGATATTATGAAGGGCCTTGAGCATCTGGATTCTG
CCTAATAAAAAACATTTATTTTCATTGCTAGCAAAAAAAAAAAAAAAAAAAAAAAAAAAAAAAAAAAA
AAAAAAAAAAAAAAAAAAAAAAAAGAAAAAAAAAAAAAAAAAAAAAAAAAAAAAAAAAAAAAAAAAAA
AAAAAAAAAAAAAAAAACGTCTTCCTGCAGGGATCCCTGGGCCTCATGGGCCTTCCGCTCACTGCCCG
CTTTCCAGTCGGGAAACCTGTCGTGCCAGCTGCATTAACATGGTCATAGCTGTTTCCTTGCGTATTGG
GCGCTCTCCGCTTCCTCGCTCACTGACTCGCTGCGCTCGGTCGTTCGGGTAAAGCCTGGGGTGCCTAA
TGAGCAAAAGGCCAGCAAAAGGCCAGGAACCGTAAAAAGGCCGCGTTGCTGGCGTTTTTCCATAGGCT
CCGCCCCCCTGACGAGCATCACAAAAATCGACGCTCAAGTCAGAGGTGGCGAAACCCGACAGGACTAT
AAAGATACCAGGCGTTTCCCCCTGGAAGCTCCCTCGTGCGCTCTCCTGTTCCGACCCTGCCGCTTACC
GGATACCTGTCCGCCTTTCTCCCTTCGGGAAGCGTGGCGCTTTCTCATAGCTCACGCTGTAGGTATCT
CAGTTCGGTGTAGGTCGTTCGCTCCAAGCTGGGCTGTGTGCACGAACCCCCCGTTCAGCCCGACCGCT
GCGCCTTATCCGGTAACTATCGTCTTGAGTCCAACCCGGTAAGACACGACTTATCGCCACTGGCAGCA
GCCACTGGTAACAGGATTAGCAGAGCGAGGTATGTAGGCGGTGCTACAGAGTTCTTGAAGTGGTGGCC
TAACTACGGCTACACTAGAAGAACAGTATTTGGTATCTGCGCTCTGCTGAAGCCAGTTACCTTCGGAA
AAAGAGTTGGTAGCTCTTGATCCGGCAAACAAACCACCGCTGGTAGCGGTGGTTTTTTTGTTTGCAAG
CAGCAGATTACGCGCAGAAAAAAAGGATCTCAAGAAGATCCTTTGATCTTTTCTACGGGGTCTGACGC
TCAGTGGAACGAAAACTCACGTTAAGGGATTTTGGTCATGAGATTATCAAAAAGGATCTTCACCTAGA
TCCTTTTAAATTAAAAATGAAGTTTTAAATCAATCTAAAGTATATATGAGTAAACTTGGTCTGACAGT
TACCAATGCTTAATCAGTGAGGCACCTATCTCAGCGATCTGTCTATTTCGTTCATCCATAGTTGCCTG
ACTCCCCGTCGTGTAGATAACTACGATACGGGAGGGCTTACCATCTGGCCCCAGTGCTGCAATGATAC

CGCGAGAACCACGCTCACCGGCTCCAGATTTATCAGCAATAAACCAGCCAGCCGGAAGGGCCGAGCGC
AGAAGTGGTCCTGCAACTTTATCCGCCTCCATCCAGTCTATTAATTGTTGCCGGGAAGCTAGAGTAAG
TAGTTCGCCAGTTAATAGTTTGCGCAACGTTGTTGCCATTGCTACAGGCATCGTGGTGTCACGCTCGT
CGTTTGGTATGGCTTCATTCAGCTCCGGTTCCCAACGATCAAGGCGAGTTACATGATCCCCCATGTTG
TGCAAAAAAGCGGTTAGCTCCTTCGGTCCTCCGATCGTTGTCAGAAGTAAGTTGGCCGCAGTGTTATC
ACTCATGGTTATGGCAGCACTGCATAATTCTCTTACTGTCATGCCATCCGTAAGATGCTTTTCTGTGA
CTGGTGAGTACTCAACCAAGTCATTCTGAGAATAGTGTATGCGGCGACCGAGTTGCTCTTGCCCGGCG
TCAATACGGGATAATACCGCGCCACATAGCAGAACTTTAAAAGTGCTCATCATTGGAAAACGTTCTTC
GGGGCGAAAACTCTCAAGGATCTTACCGCTGTTGAGATCCAGTTCGATGTAACCCACTCGTGCACCCA
ACTGATCTTCAGCATCTTTTACTTTCACCAGCGTTTCTGGGTGAGCAAAAACAGGAAGGCAAAATGCC
GCAAAAAAGGGAATAAGGGCGACACGGAAATGTTGAATACTCATACTCTTCCTTTTTCAATATTATTG
AAGCATTTATCAGGGTTATTGTCTCATGAGCGGATACATATTTGAATGTATTTAGAAAAATAAACAAA
TAGGGGTTCCGCGCACATTTCCCCGAAAAGTGCCAC

>modGLA

Cap1GGAGAACUCUUCUGGUCCCCACAGACUCAGAGAGAACCCACCGGUGCCGCCACCAUGCAGCUGA
GGAACCCAGAACUACAUCUGGGCUGCGCGCUUGCGCUUCGCUUCCUGGCCCUCGUUUCCUGGGACAUC
CCUGGGGCUAGAGCACUGGACAAUGGAUUGGCAAGGACGCCUACCAUGGGCUGGCUGCACUGGGAGCG
CUUCAUGUGCAACCUUGACUGCCAGGAAGAGCCAGAUUCCUGCAUCAGUGAGAAGCUCUUCAUGGAGA
UGGCAGAGCUCAUGGUCUCAGAAGGCUGGAAGGAUGCAGGUUAUGAGUACCUCUGCAUUGAUGACUGU
UGGAUGGCUCCCCAAAGAGAUUCAGAAGGCAGACUUCAGGCAGACCCUCAGCGCUUUCCUCAUGGGAU
UCGCCAGCUAGCUAAUUAUGUUCACAGCAAAGGACUGAAGCUAGGGAUUUAUGCAGAUGUUGGAAAUA
AAACCUGCGCAGGCUUCCCUGGGAGUUUUGGAUACUACGACAUUGAUGCCCAGACCUUUGCUGACUGG
GGAGUAGAUCUGCUAAAAUUUGAUGGUUGUUACUGUGACAGUUUGGAAAAUUUGGCAGAUGGUUAUAA
GCACAUGUCCUUGGCCCUGAAUAGGACUGGCAGAAGCAUUGUGUACUCCUGUGAGUGGCCUCUUUAUA
UGUGGCCCUUUCAAAAGCCCAAUUAUACAGAAAUCCGACAGUACUGCAAUCACUGGCGAAAUUUUGCU
GACAUUGAUGAUUCCUGGAAAAGUAUAAAGAGUAUCUUGGACUGGACAUCUUUUAACCAGGAGAGAAU
UGUUGAUGUUGCUGGACCAGGGGGUUGGAAUGACCCAGAUAUGUUAGUGAUUGGCAACUUUGGCCUCA
GCUGGAAUCAGCAAGUAACUCAGAUGGCCCUCUGGGCUAUCAUGGCUGCUCCUUUAUUCAUGUCUAAU
GACCUCCGACACAUCAGCCCUCAAGCCAAAGCUCUCCUUCAGGAUAAGGACGUAAUUGCCAUCAAUCA
GGACCCCUUGGGCAAGCAAGGGUACCAGCUUAGACAGGGAGACAACUUUGAAGUGUGGGAACGACCUC
UCUCAGGCUUAGCCUGGGCUGUAGCUAUGAUAAACCGGCAGGAGAUUGGUGGACCUCGCUCUUAUACC
AUCGCAGUUGCUUCCCUGGGUAAAGGAGUGGCCUGUAAUCCUGCCUGCUUCAUCACACAGCUCCUCCC
UGUGAAAAGGAAGCUAGGGUUCUAUGAAUGGACUUCAAGGUUAAGAAGUCACAUAAAUCCCACAGGCA
CUGUUUUGCUUCAGCUAGAAAAUACAAUGCAGAUGUCAUUAAAAGACUUACUUUAAAGCUUGCUCGCU
UUCUUGCUGUCCAAUUUCUAUUAAAGGUUCCUUUGUUCCCUAAGUCCAACUACUAAACUGGGGGAUAU
UAUGAAGGGCCUUGAGCAUCUGGAUUCUGCCUAAUAAAAAACAUUUAUUUUCAUUGCUAGCAAAAAAA
AAAAAAAAAAAAAAAAAAAAAAAAAAAAAAAAAAAAAAAAAAAAAAAAAAAAAGAAAAAAAAAAAAAA
AAAAAAAAAAAAAAAAAAAAAAAAAAAAAAAAAAAAAAAAAAAAA

>pMIVT.fLuc.BtgZI

CTAAATTGTAAGCGTTAATATTTTGTTAAAATTCGGATGTGCTGCAAGGCGAGCGGCCGCAGGCCGTC
AAGGCCGCATGGTACCTGACAGCTGTAATACGACTCACTATAAGGAGAACTCTTCTGGTCCCCACAGA
CTCAGAGAGAACCCACCGGTGCCACCATGGAAGACGCCAAAAACATAAAGAAAGGCCCGGCGCCATTC
TATCCGCTGGAAGATGGAACCGCTGGAGAGCAACTGCATAAGGCTATGAAGAGATACGCCCTGGTTCC
TGGAACAATTGCTTTTACAGATGCACATATCGAGGTGGACATCACTTACGCTGAGTACTTCGAAATGT
CCGTTCGGTTGGCAGAAGCTATGAAACGATATGGGCTGAATACAAATCACAGAATCGTCGTATGCAGT
GAAAACTCTCTTCAATTCTTTATGCCGGTGTTGGGCGCGTTATTTATCGGAGTTGCAGTTGCGCCCGC
GAACGACATTTATAATGAACGTGAATTGCTCAACAGTATGGGCATTTCGCAGCCTACCGTGGTGTTCG

TTTCCAAAAAGGGGTTGCAAAAAATTTTGAACGTGCAAAAAAAGCTCCCAATCATCCAAAAAATTATT
ATCATGGATTCTAAAACGGATTACCAGGGATTTCAGTCGATGTACACGTTCGTCACATCTCATCTACC
TCCCGGTTTTAATGAATACGATTTTGTGCCAGAGTCCTTCGATAGGGACAAGACAATTGCACTGATCA
TGAACTCCTCTGGATCTACTGGTCTGCCTAAAGGTGTCGCTCTGCCTCATAGAACTGCCTGCGTGAGA
TTCTCGCATGCCAGAGATCCTATTTTTGGCAATCAAATCATTCCGGATACTGCGATTTTAAGTGTTGT
TCCATTCCATCACGGTTTTGGAATGTTTACTACACTCGGATATTTGATATGTGGATTTCGAGTCGTCT
TAATGTATAGATTTGAAGAAGAGCTGTTTCTGAGGAGCCTTCAGGATTACAAGATTCAAAGTGCGCTG
CTGGTGCCAACCCTATTCTCCTTCTTCGCCAAAAGCACTCTGATTGACAAATACGATTTATCTAATTT
ACACGAAATTGCTTCTGGTGGCGCTCCCCTCTCTAAGGAAGTCGGGGAAGCGGTTGCCAAGAGGTTCC
ATCTGCCAGGTATCAGGCAAGGATATGGGCTCACTGAGACTACATCAGCTATTCTGATTACACCCGAG
GGGGATGATAAACCGGGCGCGGTCGGTAAAGTTGTTCCATTTTTTGAAGCGAAGGTTGTGGATCTGGA
TACCGGGAAAACGCTGGGCGTTAATCAAAGAGGCGAACTGTGTGTGAGAGGTCCTATGATTATGTCCG
GTTATGTAAACAATCCGGAAGCGACCAACGCCTTGATTGACAAGGATGGATGGCTACATTCTGGAGAC
ATAGCTTACTGGGACGAAGACGAACACTTCTTCATCGTTGACCGCCTGAAGTCTCTGATTAAGTACAA
AGGCTATCAGGTGGCTCCCGCTGAATTGGAATCCATCTTGCTCCAACACCCCAACATCTTCGACGCAG
GTGTCGCAGGTCTTCCCGACGATGACGCCGGTGAACTTCCCGCCGCCGTTGTTGTTTTGGAGCACGGA
AAGACGATGACGGAAAAAGAGATCGTGGATTACGTCGCCAGTCAAGTAACAACCGCGAAAAAGTTGCG
CGGAGGAGTTGTGTTTGTGGACGAAGTACCGAAAGGTCTTACCGGAAAACTCGACGCAAGAAAAATCA
GAGAGATCCTCATAAAGGCCAAGAAGGGCGGAAAGATCGCCGTGTAAAAGCTTGCTCGCTTTCTTGCT
GTCCAATTTCTATTAAAGGTTCCTTTGTTCCCTAAGTCCAACTACTAAACTGGGGGATATTATGAAGG
GCCTTGAGCATCTGGATTCTGCCTAATAAAAAACATTTATTTTCATTGCTAGCAAAAAAAAAAAAAAA
AAAAAAAAAAAAAAAAAAAAAAAAAAAAAAAAAAAAAAAAAAAAAGAAAAAAAAAAAAAAAAAAAAAA
AAAAAAAAAAAAAAAAAAAAAAAAAAAAAAAAAAAAAAAAGAAGAGCACGCATCGCGGATCCCTGGCC
TCATGGGCCTTCCGCTCACTGCCCGCTTTCCAGTCGGGAAACCTGTCGTGCCAGCTGCATTAACATGG
TCATAGCTGTTTCCTTGCGTATTGGGCGCTCTCCGCTTCCTCGCTCACTGACTCGCTGCGCTCGGTCG
TTCGGGTAAAGCCTGGGGTGCCTAATGAGCAAAAGGCCAGCAAAAGGCCAGGAACCGTAAAAAGGCCG
CGTTGCTGGCGTTTTTCCATAGGCTCCGCCCCCCTGACGAGCATCACAAAAATCGACGCTCAAGTCAG
AGGTGGCGAAACCCGACAGGACTATAAAGATACCAGGCGTTTCCCCCTGGAAGCTCCCTCGTGCGCTC
TCCTGTTCCGACCCTGCCGCTTACCGGATACCTGTCCGCCTTTCTCCCTTCGGGAAGCGTGGCGCTTT
CTCATAGCTCACGCTGTAGGTATCTCAGTTCGGTGTAGGTCGTTCGCTCCAAGCTGGGCTGTGTGCAC
GAACCCCCCGTTCAGCCCGACCGCTGCGCCTTATCCGGTAACTATCGTCTTGAGTCCAACCCGGTAAG
ACACGACTTATCGCCACTGGCAGCAGCCACTGGTAACAGGATTAGCAGAGCGAGGTATGTAGGCGGTG
CTACAGAGTTCTTGAAGTGGTGGCCTAACTACGGCTACACTAGAAGAACAGTATTTGGTATCTGCGCT
CTGCTGAAGCCAGTTACCTTCGGAAAAAGAGTTGGTAGCTCTTGATCCGGCAAACAAACCACCGCTGG
TAGCGGTGGTTTTTTTGTTTGCAAGCAGCAGATTACGCGCAGAAAAAAAGGATCTCAAGAAGATCCTT
TGATCTTTTCTACGGGGTCTGACGCTCAGTGGAACGAAAACTCACGTTAAGGGATTTTGGTCATGAGA
TTATCAAAAAGGATCTTCACCTAGATCCTTTTAAATTAAAAATGAAGTTTTAAATCAATCTAAAGTAT
ATATGAGTAAACTTGGTCTGACAGTTACCAATGCTTAATCAGTGAGGCACCTATCTCAGCGATCTGTC
TATTTCGTTCATCCATAGTTGCCTGACTCCCCGTCGTGTAGATAACTACGATACGGGAGGGCTTACCA
TCTGGCCCCAGTGCTGCAATGATACCGCGAGAACCACGCTCACCGGCTCCAGATTTATCAGCAATAAA
CCAGCCAGCCGGAAGGGCCGAGCGCAGAAGTGGTCCTGCAACTTTATCCGCCTCCATCCAGTCTATTA
ATTGTTGCCGGGAAGCTAGAGTAAGTAGTTCGCCAGTTAATAGTTTGCGCAACGTTGTTGCCATTGCT
ACAGGCATCGTGGTGTCACGCTCGTCGTTTGGTATGGCTTCATTCAGCTCCGGTTCCCAACGATCAAG
GCGAGTTACATGATCCCCCATGTTGTGCAAAAAAGCGGTTAGCTCCTTCGGTCCTCCGATCGTTGTCA
GAAGTAAGTTGGCCGCAGTGTTATCACTCATGGTTATGGCAGCACTGCATAATTCTCTTACTGTCATG
CCATCCGTAAGATGCTTTTCTGTGACTGGTGAGTACTCAACCAAGTCATTCTGAGAATAGTGTATGCG
GCGACCGAGTTGCTCTTGCCCGGCGTCAATACGGGATAATACCGCGCCACATAGCAGAACTTTAAAAG
TGCTCATCATTGGAAAACGTTCTTCGGGGCGAAAACTCTCAAGGATCTTACCGCTGTTGAGATCCAGT
TCGATGTAACCCACTCGTGCACCCAACTGATCTTCAGCATCTTTTACTTTCACCAGCGTTTCTGGGTG

AGCAAAAACAGGAAGGCAAAATGCCGCAAAAAAGGGAATAAGGGCGACACGGAAATGTTGAATACTCA
TACTCTTCCTTTTTCAATATTATTGAAGCATTTATCAGGGTTATTGTCTCATGAGCGGATACATATTT
GAATGTATTTAGAAAAATAAACAAATAGGGGTTCCGCGCACATTTCCCCGAAAAGTGCCAC

>modfLuc

Cap1GGAGAACUCUUCUGGUCCCCACAGACUCAGAGAGAACCCACCGGUGCCACCAUGGAAGACGCCA
AAAACAUAAAGAAAGGCCCGGCGCCAUUCUAUCCGCUGGAAGAUGGAACCGCUGGAGAGCAACUGCAU
AAGGCUAUGAAGAGAUACGCCCUGGUUCCUGGAACAAUUGCUUUUACAGAUGCACAUAUCGAGGUGGA
CAUCACUUACGCUGAGUACUUCGAAAUGUCCGUUCGGUUGGCAGAAGCUAUGAAACGAUAUGGGCUGA
AUACAAAUCACAGAAUCGUCGUAUGCAGUGAAAACUCUCUUCAAUUCUUUAUGCCGGUGUUGGGCGCG
UUAUUUAUCGGAGUUGCAGUUGCGCCCGCGAACGACAUUUAUAAUGAACGUGAAUUGCUCAACAGUAU
GGGCAUUUCGCAGCCUACCGUGGUGUUCGUUUCCAAAAAGGGGUUGCAAAAAAUUUUGAACGUGCAAA
AAAAGCUCCCAAUCAUCCAAAAAAUUAUUAUCAUGGAUUCUAAAACGGAUUACCAGGGAUUUCAGUCG
AUGUACACGUUCGUCACAUCUCAUCUACCUCCCGGUUUUAAUGAAUACGAUUUUGUGCCAGAGUCCUU
CGAUAGGGACAAGACAAUUGCACUGAUCAUGAACUCCUCUGGAUCUACUGGUCUGCCUAAAGGUGUCG
CUCUGCCUCAUAGAACUGCCUGCGUGAGAUUCUCGCAUGCCAGAGAUCCUAUUUUUGGCAAUCAAAUC
AUUCCGGAUACUGCGAUUUUAAGUGUUGUUCCAUUCCAUCACGGUUUUGGAAUGUUUACUACACUCGG
AUAUUUGAUAUGUGGAUUUCGAGUCGUCUUAAUGUAUAGAUUUGAAGAAGAGCUGUUUCUGAGGAGCC
UUCAGGAUUACAAGAUUCAAAGUGCGCUGCUGGUGCCAACCCUAUUCUCCUUCUUCGCCAAAAGCACU
CUGAUUGACAAAUACGAUUUAUCUAAUUUACACGAAAUUGCUUCUGGUGGCGCUCCCCUCUCUAAGGA
AGUCGGGGAAGCGGUUGCCAAGAGGUUCCAUCUGCCAGGUAUCAGGCAAGGAUAUGGGCUCACUGAGA
CUACAUCAGCUAUUCUGAUUACACCCGAGGGGGAUGAUAAACCGGGCGCGGUCGGUAAAGUUGUUCCA
UUUUUUGAAGCGAAGGUUGUGGAUCUGGAUACCGGGAAAACGCUGGGCGUUAAUCAAAGAGGCGAACU
GUGUGUGAGAGGUCCUAUGAUUAUGUCCGGUUAUGUAAACAAUCCGGAAGCGACCAACGCCUUGAUUG
ACAAGGAUGGAUGGCUACAUUCUGGAGACAUAGCUUACUGGGACGAAGACGAACACUUCUUCAUCGUU
GACCGCCUGAAGUCUCUGAUUAAGUACAAAGGCUAUCAGGUGGCUCCCGCUGAAUUGGAAUCCAUCUU
GCUCCAACACCCCAACAUCUUCGACGCAGGUGUCGCAGGUCUUCCCGACGAUGACGCCGGUGAACUUC
CCGCCGCCGUUGUUGUUUUGGAGCACGGAAAGACGAUGACGGAAAAAGAGAUCGUGGAUUACGUCGCC
AGUCAAGUAACAACCGCGAAAAAGUUGCGCGGAGGAGUUGUGUUUGUGGACGAAGUACCGAAAGGUCU
UACCGGAAAACUCGACGCAAGAAAAAUCAGAGAGAUCCUCAUAAAGGCCAAGAAGGGCGGAAAGAUCG
CCGUGUAAAAGCUUGCUCGCUUUCUUGCUGUCCAAUUUCUAUUAAAGGUUCCUUUGUUCCCUAAGUCC
AACUACUAAACUGGGGGAUAUUAUGAAGGGCCUUGAGCAUCUGGAUUCUGCCUAAUAAAAAACAUUUA
UUUUCAUUGCUAGCAAAAAAAAAAAAAAAAAAAAAAAAAAAAAAAAAAAAAAAAAAAAAA
AAAAAAGAAAAAAAAAAAAAAAAAAAAAAAAAAAAAAAAAAAAAAAAAAAAAAAAAAAA

>pMIVT.eGFP.BspQI

CTAAATTGTAAGCGTTAATATTTTGTTAAAATTCGGATGTGCTGCAAGGCGAGCGGCCGCAGGCCGTC
AAGGCCGCATGGTACCTGACAGCTGTAATACGACTCACTATAAGGAGAACTCTTCTGGTCCCCACAGA
CTCAGAGAGAACCCACCGGTGCCGCCACCATGGTGAGCAAGGGCGAGGAGCTGTTCACCGGGGTGGTG
CCCATCCTGGTCGAGCTGGACGGCGACGTAAACGGCCACAAGTTCAGCGTGTCCGGCGAGGGCGAGGG
CGATGCCACCTACGGCAAGCTGACCCTGAAGTTCATCTGCACCACCGGCAAGCTGCCCGTGCCCTGGC
CCACCCTCGTGACCACCCTGACCTACGGCGTGCAGTGCTTCAGCCGCTACCCCGACCACATGAAGCAG
CACGACTTCTTCAAGTCCGCCATGCCCGAAGGCTACGTCCAGGAGCGCACCATCTTCTTCAAGGACGA
CGGCAACTACAAGACCCGCGCCGAGGTGAAGTTCGAGGGCGACACCCTGGTGAACCGCATCGAGCTGA
AGGGCATCGACTTCAAGGAGGACGGCAACATCCTGGGGCACAAGCTGGAGTACAACTACAACAGCCAC
AACGTCTATATCATGGCCGACAAGCAGAAGAACGGCATCAAGGTGAACTTCAAGATCCGCCACAACAT
CGAGGACGGCAGCGTGCAGCTCGCCGACCACTACCAGCAGAACACCCCCATCGGCGACGGCCCCGTGC
TGCTGCCCGACAACCACTACCTGAGCACCCAGTCCGCCCTGAGCAAAGACCCCAACGAGAAGCGCGAT
CACATGGTCCTGCTGGAGTTCGTGACCGCCGCCGGGATCACTCTCGGCATGGACGAGCTGTACAAGTA

AGCTTGCTCGCTTTCTTGCTGTCCAATTTCTATTAAAGGTTCCTTTGTTCCCTAAGTCCAACTACTAA
ACTGGGGGATATTATGAAGGGCCTTGAGCATCTGGATTCTGCCTAATAAAAAACATTTATTTTCATTG
CTAGCAAAAAAAAAAAAAAAAAAAAAAAAAAAAAAAAAAAAAAAAAAAAAAAAAAAAAAAAAAAAGAA
AAAAAAAAAAAAAAAAAAAAAAAAAAAAAAAAAAAAAAAAAAAAAAAAAAAAAAAAAACGAAGAGCCT
GCAGTGAGGATCCCTGGGCCTCATGGGCCTTCCGCTCACTGCCCGCTTTCCAGTCGGGAAACCTGTCG
TGCCAGCTGCATTAACATGGTCATAGCTGTTTCCTTGCGTATTGGGCGCTCTCCGCTTCCTCGCTCAC
TGACTCGCTGCGCTCGGTCGTTCGGGTAAAGCCTGGGGTGCCTAATGAGCAAAAGGCCAGCAAAAGGC
CAGGAACCGTAAAAAGGCCGCGTTGCTGGCGTTTTTCCATAGGCTCCGCCCCCCTGACGAGCATCACA
AAAATCGACGCTCAAGTCAGAGGTGGCGAAACCCGACAGGACTATAAAGATACCAGGCGTTTCCCCCT
GGAAGCTCCCTCGTGCGCTCTCCTGTTCCGACCCTGCCGCTTACCGGATACCTGTCCGCCTTTCTCCC
TTCGGGAAGCGTGGCGCTTTCTCATAGCTCACGCTGTAGGTATCTCAGTTCGGTGTAGGTCGTTCGCT
CCAAGCTGGGCTGTGTGCACGAACCCCCCGTTCAGCCCGACCGCTGCGCCTTATCCGGTAACTATCGT
CTTGAGTCCAACCCGGTAAGACACGACTTATCGCCACTGGCAGCAGCCACTGGTAACAGGATTAGCAG
AGCGAGGTATGTAGGCGGTGCTACAGAGTTCTTGAAGTGGTGGCCTAACTACGGCTACACTAGAAGAA
CAGTATTTGGTATCTGCGCTCTGCTGAAGCCAGTTACCTTCGGAAAAAGAGTTGGTAGCTCTTGATCC
GGCAAACAAACCACCGCTGGTAGCGGTGGTTTTTTTGTTTGCAAGCAGCAGATTACGCGCAGAAAAAA
AGGATCTCAAGAAGATCCTTTGATCTTTTCTACGGGGTCTGACGCTCAGTGGAACGAAAACTCACGTT
AAGGGATTTTGGTCATGAGATTATCAAAAAGGATCTTCACCTAGATCCTTTTAAATTAAAAATGAAGT
TTTAAATCAATCTAAAGTATATATGAGTAAACTTGGTCTGACAGTTACCAATGCTTAATCAGTGAGGC
ACCTATCTCAGCGATCTGTCTATTTCGTTCATCCATAGTTGCCTGACTCCCCGTCGTGTAGATAACTA
CGATACGGGAGGGCTTACCATCTGGCCCCAGTGCTGCAATGATACCGCGAGAACCACGCTCACCGGCT
CCAGATTTATCAGCAATAAACCAGCCAGCCGGAAGGGCCGAGCGCAGAAGTGGTCCTGCAACTTTATC
CGCCTCCATCCAGTCTATTAATTGTTGCCGGGAAGCTAGAGTAAGTAGTTCGCCAGTTAATAGTTTGC
GCAACGTTGTTGCCATTGCTACAGGCATCGTGGTGTCACGCTCGTCGTTTGGTATGGCTTCATTCAGC
TCCGGTTCCCAACGATCAAGGCGAGTTACATGATCCCCCATGTTGTGCAAAAAAGCGGTTAGCTCCTT
CGGTCCTCCGATCGTTGTCAGAAGTAAGTTGGCCGCAGTGTTATCACTCATGGTTATGGCAGCACTGC
ATAATTCTCTTACTGTCATGCCATCCGTAAGATGCTTTTCTGTGACTGGTGAGTACTCAACCAAGTCA
TTCTGAGAATAGTGTATGCGGCGACCGAGTTGCTCTTGCCCGGCGTCAATACGGGATAATACCGCGCC
ACATAGCAGAACTTTAAAAGTGCTCATCATTGGAAAACGTTCTTCGGGGCGAAAACTCTCAAGGATCT
TACCGCTGTTGAGATCCAGTTCGATGTAACCCACTCGTGCACCCAACTGATCTTCAGCATCTTTTACT
TTCACCAGCGTTTCTGGGTGAGCAAAAACAGGAAGGCAAAATGCCGCAAAAAAGGGAATAAGGGCGAC
ACGGAAATGTTGAATACTCATACTCTTCCTTTTTCAATATTATTGAAGCATTTATCAGGGTTATTGTC
TCATGAGCGGATACATATTTGAATGTATTTAGAAAAATAAACAAATAGGGGTTCCGCGCACATTTCCC
CGAAAAGTGCCAC

>modEGFP

Cap1GGAGAACUCUUCUGGUCCCCACAGACUCAGAGAGAACCCACCGGUGCCGCCACCAUGGUGAGCA
AGGGCGAGGAGCUGUUCACCGGGGUGGUGCCCAUCCUGGUCGAGCUGGACGGCGACGUAAACGGCCAC
AAGUUCAGCGUGUCCGGCGAGGGCGAGGGCGAUGCCACCUACGGCAAGCUGACCCUGAAGUUCAUCUG
CACCACCGGCAAGCUGCCCGUGCCCUGGCCCACCCUCGUGACCACCCUGACCUACGGCGUGCAGUGCU
UCAGCCGCUACCCCGACCACAUGAAGCAGCACGACUUCUUCAAGUCCGCCAUGCCCGAAGGCUACGUC
CAGGAGCGCACCAUCUUCUUCAAGGACGACGGCAACUACAAGACCCGCGCCGAGGUGAAGUUCGAGGG
CGACACCCUGGUGAACCGCAUCGAGCUGAAGGGCAUCGACUUCAAGGAGGACGGCAACAUCCUGGGGC
ACAAGCUGGAGUACAACUACAACAGCCACAACGUCUAUAUCAUGGCCGACAAGCAGAAGAACGGCAUC
AAGGUGAACUUCAAGAUCCGCCACAACAUCGAGGACGGCAGCGUGCAGCUCGCCGACCACUACCAGCA
GAACACCCCCAUCGGCGACGGCCCCGUGCUGCUGCCCGACAACCACUACCUGAGCACCCAGUCCGCCC
UGAGCAAAGACCCCAACGAGAAGCGCGAUCACAUGGUCCUGCUGGAGUUCGUGACCGCCGCCGGGAUC
ACUCUCGGCAUGGACGAGCUGUACAAGUAAGCUUGCUCGCUUUCUUGCUGUCCAAUUUCUAUUAAAGG
UUCCUUUGUUCCCUAAGUCCAACUACUAAACUGGGGGAUAUUAUGAAGGGCCUUGAGCAUCUGGAUUC

UGCCUAAUAAAAAACAUUUAUUUUCAUUGCUAGCAAAAAAAAAAAAAAAAAAAAAAAAAAAAAAAAAA
AAAAAAAAAAAAAAAAAAAAAAAAAAGAAAAAAAAAAAAAAAAAAAAAAAAAAAAAAAAAAAAAAAAAAAAAAAAAA
AAAAAAAAAAAAAAAAAA
